# High Throughput Hyperspectral and Multiplexed Super-Resolution Fluorescence Imaging by SP-STORM

**DOI:** 10.1101/2025.09.17.676812

**Authors:** Elric Dion Pott, Meek Yang, James Ethan Batey, Joie Embree, Bin Dong

## Abstract

Simultaneous determination of spatial location and spectral color of single molecules at large molecular density with high throughput was achieved by combining single molecule photoswitching and in-hardware transformation based spectral phasor analysis. The method, named spectral phasor enabled stochastic optical reconstruction microscopy (SP-STORM), achieved simultaneous super-resolution imaging of five subcellular structures with minimum crosstalk for the first time. The high throughput feature of SP-STORM enables these subcellular structures to be readily resolved in about one minute, which is more than a magnitude faster than other multiplexing single molecule localization microscopy techniques. The concept of SP-STORM is also compatible with and can be readily applicable to other super-resolution microscopy.

## Main

In the past decades, the development of single molecule localization based super-resolved fluorescence microscopy (SMLM) ^1, 2^ enables studies of protein assemblies down to nanometer spatial resolution and resolves the spatial arrangements of protein organization at single-protein resolution.^3, 4^ Further advancement in multicolor SMLM attempts to explore its multiplexing capability. Utilizing spectrally well-separated fluorophores^5, 6^ is the most straightforward approach, yet it requires long data acquisition time due to sequential image registration for each color and involves complex error-prone alignment procedures between channels. Therefore, its throughput is low. Ratiometric-based spectral demixing approach enables simultaneously resolving up to four targets but suffers high crosstalk.^7^ Dispersive (i.e., SR-STORM^8^) and excitation modulation (i.e., ExR-STORM^9^) based spectral demixing approaches have pushed the multiplexing capability to simultaneous determination of four targets with low crosstalk. However, both suffers low throughput due to requiring low molecular density to avoid spectral interference (SR-STORM) and needing mechanically switching between excitation laser lines (ExR-STORM). Four-color SR-STORM and ExR-STORM take12 and 25 minutes respectively. In comparison, single-color dSTORM requires less than a minute.^10, 11^

DNA-PAINT is another type of multiplexing SMLM with high spatial resolution.^12^ It has theoretically-unlimited multiplexing capability.^13^ However, the throughput of this approach remains a major limitation. Recent optimizations, such as FRET-based probes^14^ and fluorogenic DNA-PAINT,^15^ have improved acquisition speed at the expense of reducing multiplexing capacity. The development of secondary label-based unlimited multiplexed PAINT (SUM-PAINT)^16^ attempted to address this trade-off by decoupling the DNA barcoding of the target from the imaging process. The time to quantitative map up to 30 proteins in neuron cells at single-protein resolution was reduced from 800 hours using Exchange-PAINT^13^ to 30 hours using SUM-PAINT. Nevertheless, the intrinsic nature of sequential imaging in SUM-PAINT remains as the bottleneck for high throughput analysis.

Here, we developed an approach to simultaneously determine single-molecule locations and their fluorescence emission spectra with high throughput. The approach is based on in-hardware spectral phasor analysis. We simultaneously obtained fluorescence images of single molecules with and without their emission modulated by wavefront-like transmission optical filters. The spectral color of single molecules was directly determined from their relative photon intensities with and without modulation. This approach is non-dispersive and does not require iterative switching between channels thus eliminating spectral interferences, improving signal to noise ratio, and allowing image acquisition at high speed and large molecule density. Combining single molecule photoswitching, high throughput hyperspectral and multiplexed super-resolution microscopy imaging was achieved. In this work, we integrate the in-hardware spectral phasor analysis with dSTORM^10^ imaging, named as SP-STORM. We obtained simultaneously multiplexed super-resolution microscopy image of 5 protein targets in ∼1 min. This is more than a magnitude order faster than all previously developed multiplexing SMLM techniques which can only simultaneously resolve up to 4 protein targets in few tens of minutes to hours.

To realize SP-STORM, we modulated single fluorescent molecules’ emission using a lab-built three-channel imager (Fig. 1a, Supplementary Fig. 1), where the signals in two of the channels were transformed by wavefront-like transmission optical filters (Fig. 1a, insert), namely the sine and cosine channels. The unmodified signal in the remaining channel works as the reference for the phasor analysis. From photon intensities of the same single molecules in three channels, one can obtain their locations in the phasor space (Supplementary 2) where the phase angle (ϕ) is wavelength dependent. Ideally, the optical filters should have transmission profiles representing sine and cosine functions.^17, 18^ However, optical filters with broadband transmission similar to ideal sine/cosine filters can achieve the same purpose with modified analysis procedure (Supplementary 3).^19^ We adapted two optical filters with their transmission profiles mimicking half of the sine and cosine functions in wavelength range from 635 to 750 nm (Fig. 1a, insert), matching the spectral emission of far-red dyes for dSTORM imaging. To determine the spectral color of single molecules in our imaging setup, we calibrated the imaging system by refining the spectral color of light from microscope built-in lamp or emission from fluorescence beads using narrow bandpass filters (Fig. 1b-c). Photon intensities of lamp or same fluorescent beads in three channels were measured and used to calculate their locations the phasor space (Fig.1d). The spectral mean-phase angle relationship was obtained from the phasor plot and follows a nonlinear trend that was well-fitted by a fourth-order polynomial function (Fig. 1e). This calibration curve was then used to determine the spectral mean of single molecules in dSTORM imaging experiments.

**Figure 1.**
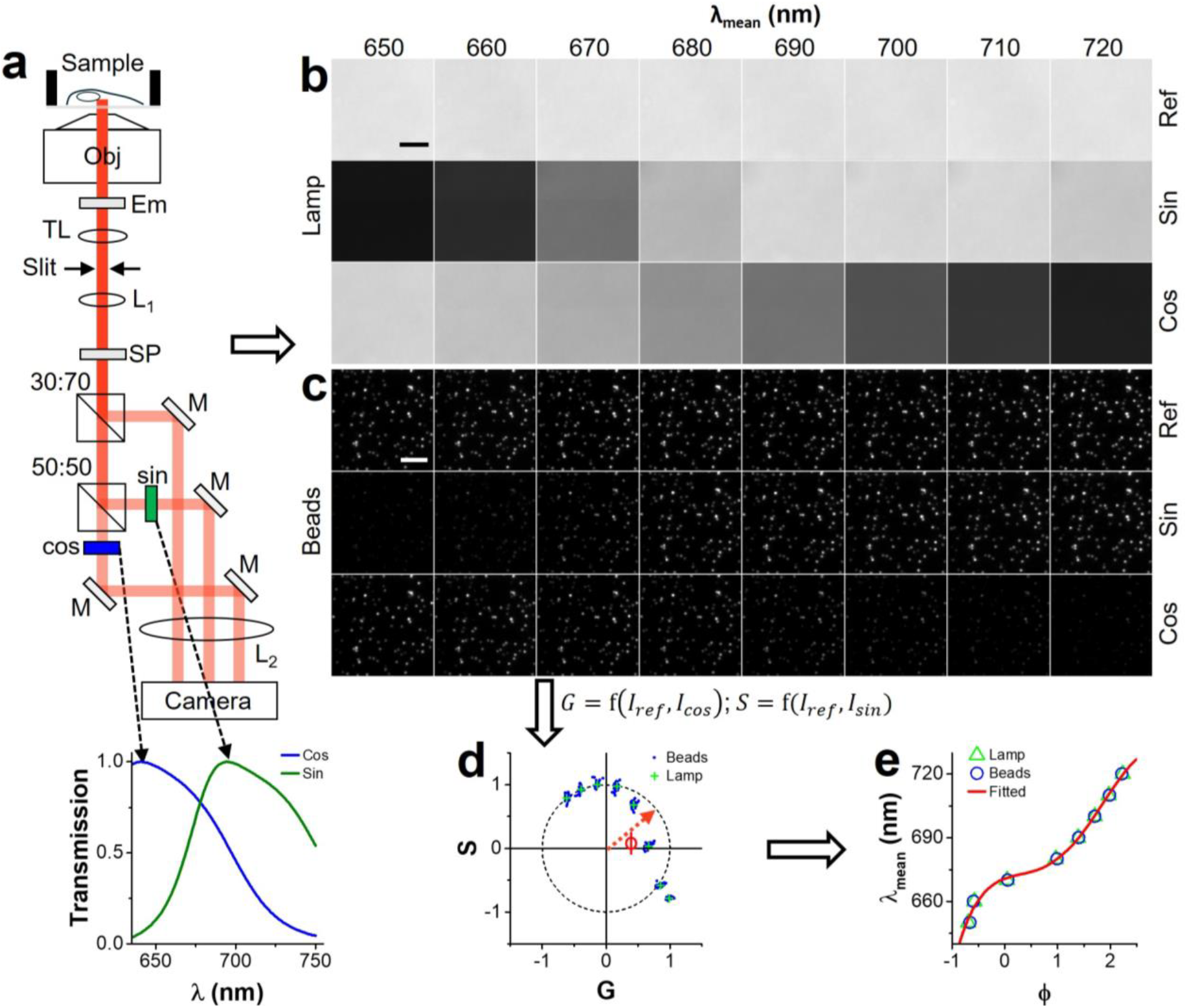
Working principle of SP-STORM. (a) Schematic of the optical setup. Collected emission signals from single molecules are separated into three channels with two of them transformed by sine and cosine optical filters. DC, dichroic mirror; Em, long pass emission filter; TL, tube lens; L, lens; SP, short pass filter; M, mirror. Insert: Normalized transmission profiles of ideal and actual sine and cosine filters in confined wavelength range using the Em and SP filters. (b) Brightfield image of microscope lamp light and (c) fluorescence image of beads at different wavelengths. (d) Phasor plot of results at different wavelengths. (e) The wavelength-phase angle relationship for system calibration. The data was fitted by a fourth-order polynomial function. Scale bar: 5 µm.

We first tested one of the most widely used dSTORM dye, Alexa fluor 647 (AF647) for SP-STORM imaging. We labelled the fixed cells with the AF647 dye through immunostaining and then photoswitched most of the dye molecules into a non-fluorescent dark state (i.e., off state) resulting in only a subset of dye molecules fluorescing (i.e., on state) at any given instant. By optimizing the excitation laser power density (>10 kW cm^-2^) and imaging acquisition rate (200 Hz), the transformed (i.e., cosine and sine) and unmodified (i.e., reference) emission signal from sparsely distributed single AF647 molecules were simultaneously recorded on different regions of the same camera (Fig. 2a). Single AF647 molecules’ positions were first localized. Same single molecules in three channels were identified and their weighted locations were used for image reconstruction. The photon intensities of the same single AF647 molecules in three channels were measured and used to determine their locations and phase angles in the phasor space (Fig. 2b). The spectral means of individual molecules were then determined from the measured phase angles by using the established calibration curve (Fig. 1e). The targeted subcellular structures were reconstructed using super-localized positions of single AF647 molecules with color representing by the determined spectral mean of single molecules ranging from 660-720 nm (Fig 2c). The localization precision for AF647 was estimated to be ∼6 nm from small cluster analysis (Supplementary Fig. 7), which corresponds to a spatial resolution of ∼15 nm (full width at half maximum) comparable to typical dSTORM imaging. The histogram distribution of > 10^5^ single AF647 molecules (Fig. 2d) gives an average spectral mean of 674.2 nm and a spectral variation of 1.4 nm (s.d.).

**Figure 2.**
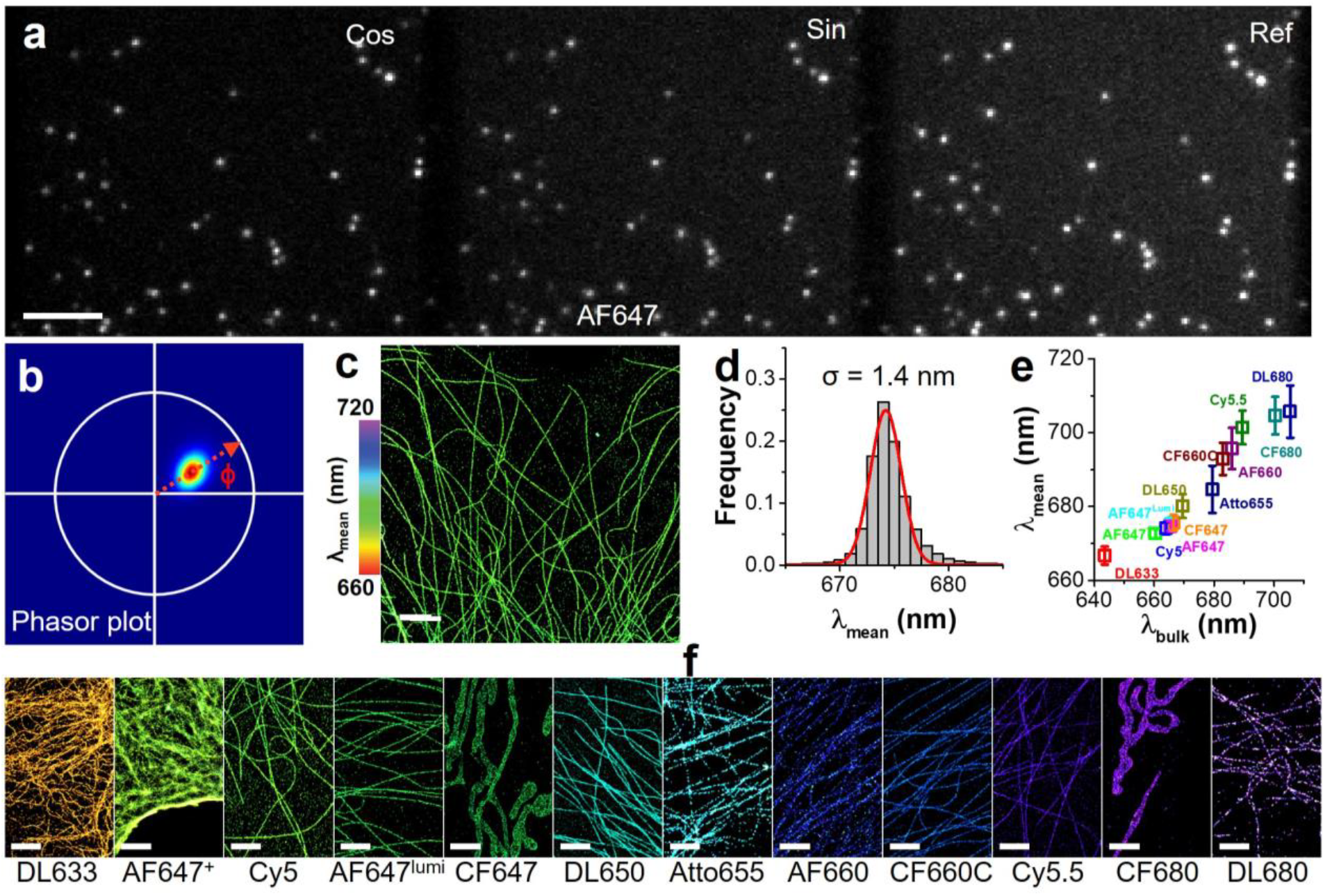
SP-STORM imaging with 13 far-red dyes. (a) Simultaneous acquiring of transformed (i.e., sine/cosine) and unmodified (i.e., ref) images from the same single AF647 molecules. (b) Phasor plot of >10^5^ single AF647 molecules. (c) Super-resolution image of labeled microtubule with color representing single-molecule spectral mean. (d) 1D Gaussian fitting the histogram of the spectral mean of single AF647 molecules gives an average of 674.2 ± 1.4 nm (mean ± s.d.) (e) Measured spectral mean for single molecules of 13 tested far-red dyes. Error bars are s.d. from Gaussian fitting of the histogram distribution of spectral mean from single molecules (> 10^5^ for each dye). (f) Super-resolution image of labeled subcellular structures for different dyes with color representing their single-molecule spectral mean. Scale bars: (a) 5 µm, (c, f) 2 µm.

The small spectral variation holds promises for simultaneously multiplexed SMLM by spectral demixing dyes with minimal spectral difference. To test the feasibility of this concept, we tested another 12 far-red organic dyes using SP-STORM with the same imaging setup as testing AF647 (Supplementary Fig. 4-16). The spatial resolution ranges from 15-40 nm. The results of spectral mean and variation over > 10^5^ single molecules for each dye are shown in Fig. 2e. The spectral variation ranges from 1.3-7.1 nm. Representing super-resolution images with color representing spectral means of single molecules for these 12 far-red dyes were shown in Fig. 2f. Among these dyes, AF plus 647 and DL680 have the smallest and largest spectral variation, respectively. Interestingly, we found out most of the tested dyes show a single population of spectral color, except the AF660 and DL680 (Supplementary Fig. 12, 16). These results laid the foundation for multiplexed SMLM by choosing dyes based on photoswitching capability, spatial resolution, spectral variation, and overlapping in the phasor space.

To test SP-STORM for multiplexed imaging and compare the throughput to previously published works, we labelled four distinct subcellular targets in fixed cells with four dyes (i.e., DL633, AF647, CF660C, and CF680) that have heavily overlapped spectra. A single red laser was used for exciting and photoswitching all dye molecules between on and off states (Fig. 3a). The targeted subcellular structures were reconstructed using super-localized positions of single molecules with color representing by the spectral mean of single molecules ranging from 660-720 nm (Fig. 3c, Supplementary Fig. 17a). Excitingly, the targeted four subcellular structures are readily distinguishable based on the spectral mean alone. Distinct colors of purple, blue, green and yellow were clearly observed for labelled mitochondria, microtubule, vimentin, and peroxisome respectively. Single molecules were further classified by comparing obtained single molecules’ location in the phasor space to that of known dye molecules (Supplementary Fig. 17g). dSTORM images of labelled subcellular structures show negligible misidentification (Fig. 3e-i, Supplementary Fig. 17b-f) with color crosstalk < 2% (Fig. 3j). We then evaluated the throughput of SP-STORM. The first and last frames of imaging data are shown in Fig. 3a. Clearly, the molecular density is large. The number of locations per image frame along with the data acquisition time was plotted in Fig. 3b. The average number of localizations per image frame was estimated to be ∼74, corresponding a density of ∼ 0.14 locations per µm^2^. Under this density of single molecules, multiplexed dSTORM images with highly resolvable subcellular structures are already readily obtained less than a minute (Fig. 3d) based on the Nyquist limit^20^ (Supplementary Fig. 18). In comparison, other simultaneously multiplexed SMLM requires few tens of minutes to hours to resolve four subcellular structures.

**Figure 3.**
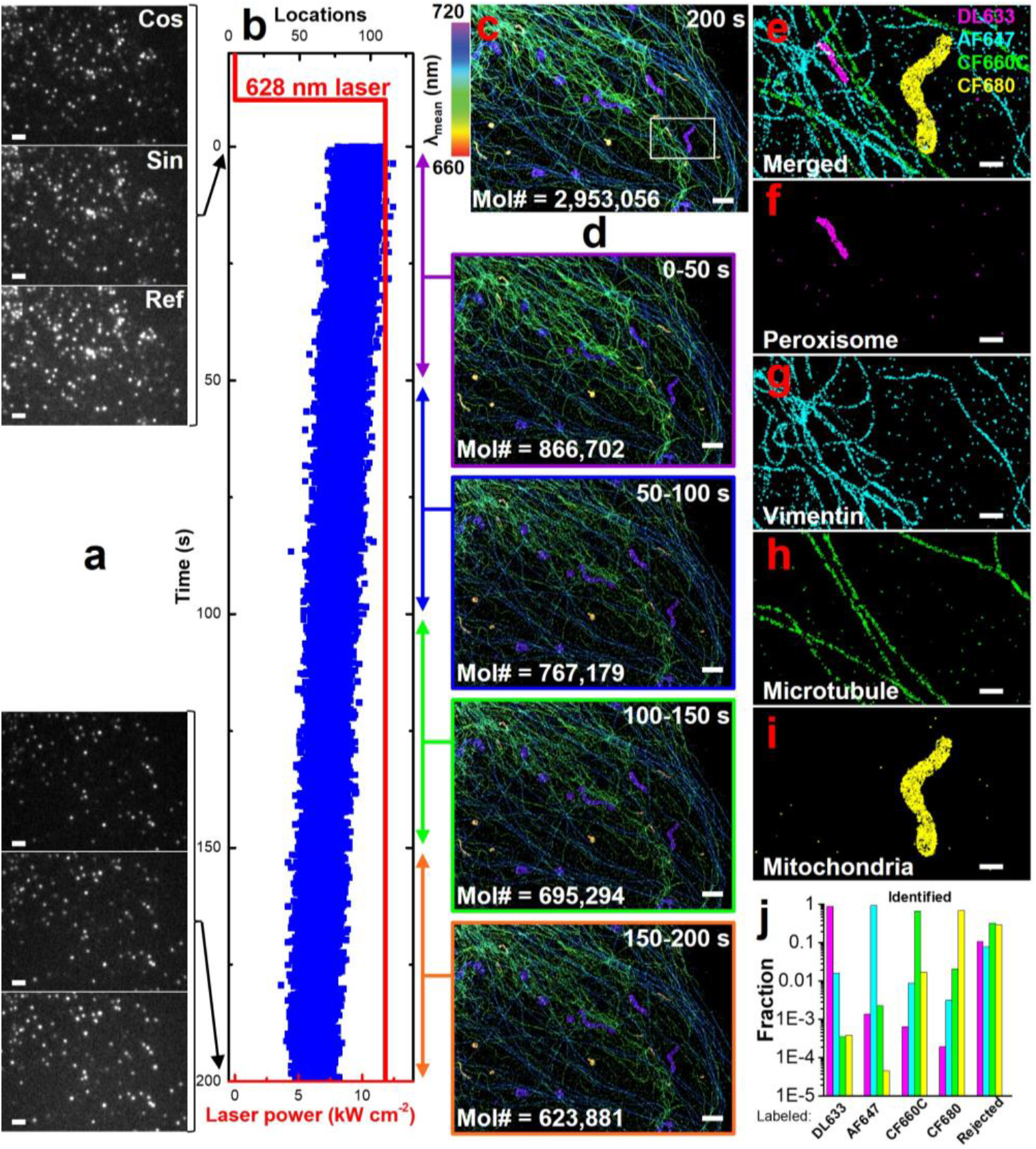
SP-STORM enables hyperspectral and simultaneously multiplexed imaging with high throughput. (a) First and last image frame of single molecules in four-color SP-STORM. (b) Trajectory of number of locations and laser power density versus time. (c) Hyperspectral dSTORM image of four proteins (i.e., PMP70, vimentin, α-tubulin, TOM20) labelled by DL633, AF647, CF660C, and CF680 in fixed COS-7 cells. The color denotes the spectral mean of single molecules. (d) Hyperspectral dSTORM image for time segments of 50 seconds achieving minimal density based on the Nyquist criterion. Mol# represents total number of molecular locations obtained for image reconstruction. (e-i) dSTORM images of four proteins after classification of single molecules. (j) Color crosstalk between channels. Scale bars: 2 µm (a, c-d) and 500 nm (e-i).

We further tested the feasibility of SP-STORM for simultaneously five-color super-resolution imaging. We labelled five distinct subcellular targets in fixed cells with five chosen dyes (i.e., DL633, AF plus 647, DL650, CF660C, and CF680) based on their spectral mean and variation determined previously (Fig. 2e). Five subcellular structures are readily observed with distinct colors (Supplementary Fig. 4a), where purple, blue, cyan, yellow-green, and yellow denotes mitochondria, microtubule, vimentin, actin, and peroxisome correspondingly. The multiplexed dSTORM image was readily obtained in ∼1.3 minutes (Supplementary Fig. 19). For all channels, the crosstalk was determined to be < 5% (Fig. 4b-i). Simultaneous 5-color SMLM imaging has never been achieved previously. To evaluate if the super-localized positions align well for different dye molecules, we performed SP-STORM imaging of mitochondrial outer membrane protein (i.e., TOM20) and mitochondrial heat shock protein (i.e., mtHsp70 or Mortalin, primarily localizes inside the mitochondria) with CF680 and DL650. The hyperspectral and multiplexed dSTORM images (Fig. 4j, k) and the cross-section profile (Fig. 4l) clearly match the expectation that locations of DL650 are fully surrounded by CF680.

**Figure 4.**
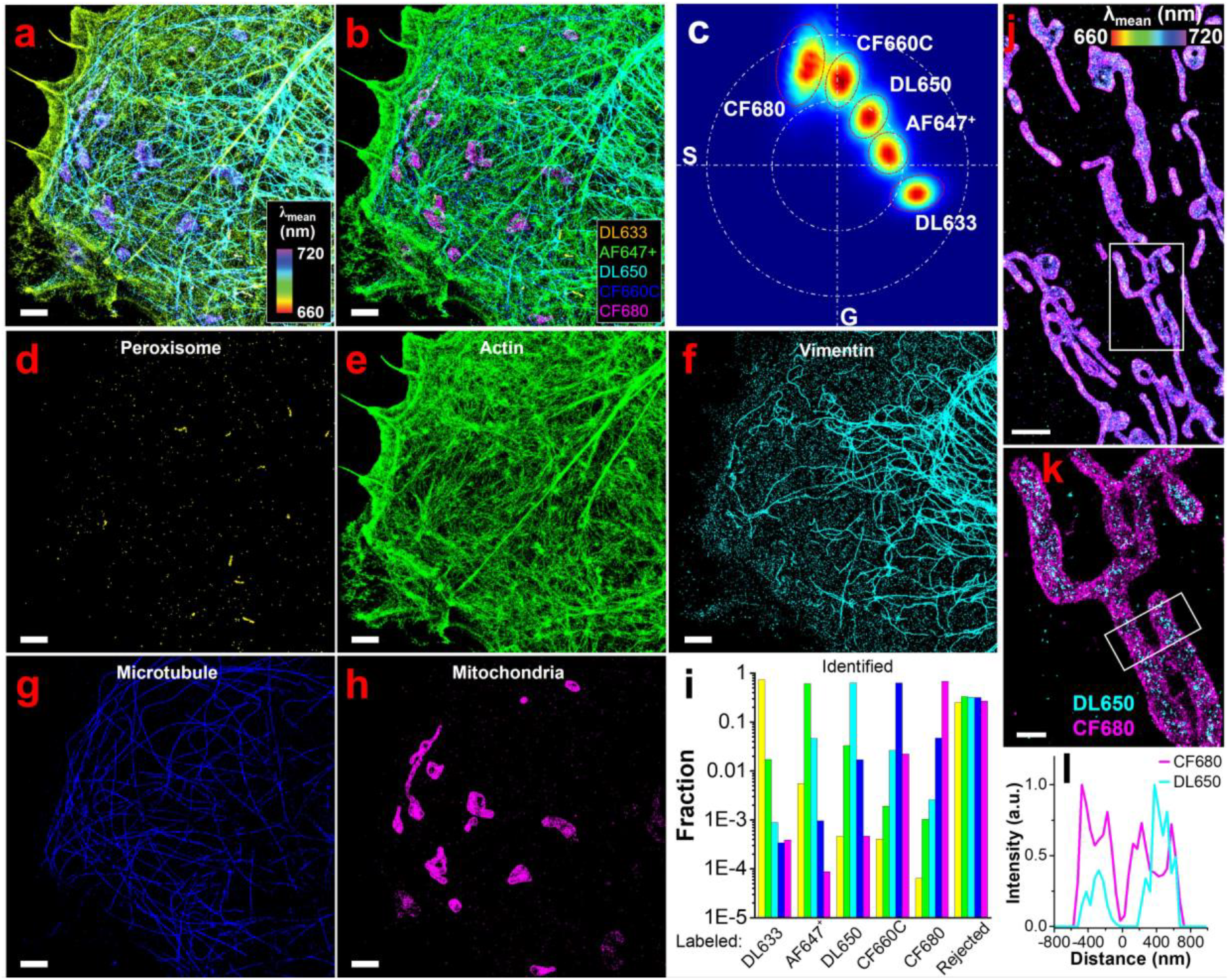
5-color SP-STORM. (a) Hyperspectral dSTORM image of five proteins (i.e., PMP70, F-actin, vimentin, α-tubulin, TOM20) labelled by DL633, AF647^+^, DL650, CF660C, and CF680 in fixed COS-7 cells. (b) 5-color dSTORM image with single molecules being classified and recolored. The classification was based on the comparison of single molecules’ location in phasor space to five dyes (c). Boundary conditions in phasor plot were established from single-color labelled sample and used to separate the different dye molecules. The red dash dot line denotes 1.2-2.0 s.d. of distributions for each dye. The white dash dot line is for illustration location of dyes in the phasor plot with large and small circles as unit and half-unit circle respectively. (d-h) The separated channels of five subcellular structures. (i) Color crosstalk between channels. (j) Hyperspectral and (k) 2-color dSTORM images (white box in (j)) of TOM20 and mortalin labelled with CF680 and DL650 respectively. (l) Cross-section profile of two dyes along the long axis of white box in (k). Scale bars: 2 µm (a-h, j) and 500 nm (k).

The key challenges in advancing techniques for quantifying biomolecules and their spatial locations include sensitivity, spatial resolution, multiplexing capability, and throughput. Previously, the advancement of SMLM techniques has enabled the simultaneous measurement of up to four protein targets or sequential quantification of theoretically unlimited protein targets with spatial resolution at the scale matching the size of proteins. However, the intrinsic nature of these techniques, either low SNR, low molecular density, low data acquisition rate, or sequential data acquisition, limits their throughput. The unique strength of SP-STORM lies in its ability to simultaneously determine the spatial locations and spectral color of single molecules using non-dispersive approach and broadband wavefront-like transmission optical filters. It enables high-density single-molecule imaging with high SNR, resulting in both high throughput and large multiplexing capability. SP-STORM has demonstrated the capacity to resolve five distinct protein targets simultaneously and with low crosstalk in about a minute, which is more than a magnitude order faster than all current multiplexed SMLM techniques (i.e., a few tens of minutes to hours for four protein targets). While current demonstrations focused on 2D imaging, SP-STORM is readily extendable to 3D dSTORM imaging. The high throughput feature of SP-STORM also holds promises for studying dynamic processes in live cells. Furthermore, the integration of spectral phasor analysis for spectral demixing of fluorophores is broadly compatible with other super-resolution techniques, e.g., (F)PALM, PAINT, and STED, highlighting its versatility and potential for advancing multiplexed microscopy across various imaging platforms.

## Methods

### Optical setup

The SP-STORM was carried out on an Olympus IX-81 inverted microscope (Supplementary Fig. 1) equipped with an oil immersion objective (UPLAPO100X, NA 1.50, Olympus). 405 nm (Coherent) laser was coupled into an optical fiber, recollimated by a fiber coupler, and focused on the back focal plane of the oil immersion objective by a lens (L5) after passing a pair of relay lens (L3, L4). Laser at 628 nm (MPB Communications) was combined into the same optical path of 405 nm laser line using a short pass dichroic mirror (DC1) after passing a pair of relay lens (L1, L2). Lasers were directed into the sample by a dichroic mirror (DC2). All Lasers were cleaned by narrow bandpass filters (Ex1 and Ex2). Two translation stages were used for shifting the laser beams laterally before entering the objective so that the laser beams project to the sample/coverslip interface at an angle slightly smaller than the critical angle. The emission signals from dye molecules were collected by the same oil immersion objective. A long pass emission filter (RET638lp, Chroma) was used to reject laser scattering background. Then, the collected signal was directed to a lab-built three-channel imager for in-hardware transformation based spectral phasor analysis. Inside the three-channel imager, the emission signal from dye molecules was first split by a 30:70 (R:T) nonpolarized beam splitter (BS019, Thorlabs). The transmitted light after the first beam splitter was split further by a 50:50 (R:T) nonpolarized beam splitter (BS013, Thorlabs). The reflected and transmitted signals from the second beam splitter were transformed by sine and cosine function-like optical filters respectively. Three images of the same dye molecules, i.e., reference, sin/cos-modified images, were projected on different regions of the same electron-multiplying charge-coupled device (EMCCD) camera (iXon Ultra 897, Andor) using a pair of relay lens (L6, L7). A short pass optical filter (FESH0750, Thorlabs) was used to confined spectral window below 750 nm. An optical slit was placed at the intermediate image plane of tube lens allowing crop field of view and avoiding crosstalk between three channels. The sample stage was locked by a self-built autofocusing optical system running by self-written Python scripts.

### Spectral calibration

We applied two approaches to calibrate the wavelength-phase angle relationships of the SP-STORM imaging system. In one approach, brightfield images of transmitted light from the microscope lamp were captured. The lamp light was filtered by a series of narrow bandpass filters (Thorlabs) giving three images (i.e., reference, sine-, and cosine-modified) at specific wavelengths (Fig. 1b). Measuring the intensities from three channels allows the determination of locations of spectral color in the phasor plot (Fig. 1d). In another approach, 40 nm dark red fluorescent beads (F8789, ThermoFisher) were drop-casted on a coverslip at low density. The sample loaded with fluorescent beads was mounted on the microscope and excited by the lasers at a weak intensity. Similarly, narrow bandpass filters (Thorlabs) were inserted in the detection optical pathways. Three images (i.e., reference, sine-, and cosine-modified) of the fluorescent emission from the same single beads at specific wavelengths were captured (Fig. 1c). The locations and photon intensities of single fluorescent beads in all three channels were super-localized. The same fluorescent beads in three channels were identified through previous developed image projection procedure.^21^ The location of specific wavelength in the phasor plot was calculated by the photon intensities of beads in three channels (Fig. 1d). Results from both methods generated similar calibration curves that were well fitted by a fourth-order polynomial function (Fig. 1e).

### Dyes and antibodies

These dyes were obtained from ThermoFisher (DyLight633, DyLight650, DyLight680, Alexa Fluor 647, and Alexa Fluor 660) as gifts and purchased from Lumiprobe (Alexa Fluor 647 and Cyanine5), Sigma (Atto655), Biotium (CF647, CF660C, and CF680), and Cytiva (Cy5.5). All these dyes were in the form of N-hydroxysuccinimidyl ester (NHS) for conjugation to secondary antibodies. The dye-to-antibody ratio was controlled to be less than one. Primary antibodies were obtained from ThermoFisher (rat antibody to alpha tubulin, MA1-80017; mouse antibody to beta tubulin, MA5-16308; goat antibody to HSPA9, PA5-48035; rabbit antibody to TOM20, MA5-34964) as gifts and purchased from Sigma (Chicken antibody to vimentin, AB5733; mouse antibody to PMP70, SAB4200181). Unconjugated secondary antibodies (Donkey anti-rat, 712-005-153; Donkey anti-mouse, 715-005-151; Donkey anti-rabbit, 711-005-152; Bovine anti-goat, 805-005-180) were purchased from Jackson ImmunoResearch Laboratories. For labelling actin, Alexa Fluor Plus 647 phalloidin were obtained from ThermoFisher as a gift. Nile red was purchased from ThermoFisher (415711000).

### Sample preparation

#### Cell culture

COS-7, African green monkey kidney cells (CRL-1651, ATCC) was purchased from the American Type Culture Collection (ATCC). The COS-7 cells were cultured in T25 cell culture flask (690160, Greiner Bio-One) with the cell culture medium Dulbecco’s Modified Eagle’s Medium (DMEM) (10-014-CV, Corning) added with 10% fetal bovine serum (FBS) (26140079, Gibco) along with 1% Penicillin-Streptomycin (Pen Strep) (15140122, Gibco) (referred as the complete cell culture media in the rest of the article). µ-slide 8 Well high Glass Bottom chambered coverslip (80807, ibidi) was purchased from ibidi for subculturing cells. The chambers were rinsed with 1×PBS (10010023, Gibco) to remove glass dust before use. To subculture cells, to each well, 30 µL cell suspension solution was added. Then, 270 µL of complete cell culture media was added to every well. The chambered coverslip was kept in the cell culture incubator at 37°C with 5% CO_2_ for 24 hours before the cells were used in imaging experiments. µ-dish (81158, ibidi) were also used in the imaging experiments. To subculture cells in µ-dish, 120 µL of cell suspension solution was added into the dish, followed by 2 mL of complete cell culture media. The cells were allowed to incubate for 24 hours at 37°C and 5% CO_2_ inside the cell culture incubator before imaging.

#### Cell labelling with far-red dyes

After 24 hours, COS-7 cells were washed once with pre-warmed 1×PBS buffer (10010049, ThermoFisher), fixed with pre-warmed 3% paraformaldehyde (0433689M, ThermoFisher) + 0.1% glutaraldehyde (340855, Sigma) in 1×PBS buffer for 10 minutes, reduced with freshly prepared 0.3% NaBH_4_ (BDH4604, VWR) for 10 minutes, followed by five-time washes with 1×PBS buffer. The cells were blocked and permeabilized in blocking buffer containing 3% BSA (001-000-162, Jackson ImmunoResearch Laboratories), 0.2% Triton-X100 (807426, MP Biomedicals) in 1×PBS buffer for 2 hours, followed by incubation with primary antibodies in blocking buffer at 4 °C overnight. After primary antibody incubation, the cells were washed five times with washing buffer (0.75% BSA, 0.05% Triton-x100 in 1×PBS buffer), followed by incubation with dye-conjugated secondary antibodies in blocking buffer at room temperature for 2 hours. After secondary antibody incubation, the cells were washed five times with washing buffer, washed once with 1×PBS buffer, post-fixed with 3% paraformaldehyde and 0.1% glutaraldehyde in 1×PBS buffer for 10 minutes, followed by three-time wash with 1×PBS buffer. The cell sample was stored in 1×PBS buffer before microscopy imaging. For long-term storage, 20 mM (0.1%) sodium azide (14314.22, ThermoFisher) was added into the 1×PBS buffer. To co-stain actin with other subcellular structures, Alexa Fluor Plus 647 phalloidin (0.1 µM in 1×PBS buffer) was added to the cells after the post-fixation step, followed by incubation at 4 °C overnight. The cells were briefly washed two times with 1×PBS buffer before imaging.

### SP-STORM imaging

The dye labelled samples were first excited at low laser power for locating cells and capturing conventional fluorescence images. The laser power was then increased to high power density (628 nm, ∼ 12 kW cm^-2^) to photoswitch most of the dye molecules into nonfluorescent dark state. A random subset and spatially resolved dye molecules were kept in fluorescing (on state) at any given instant. 405 nm laser at low power density was used to adjust the density of single dye molecules per image frame when it is necessary. The EMCCD camera acquired images in all three channels (i.e., reference, sine- and cosine-modified) simultaneously and continuously at a frame rate of 200 Hz. The imaging data was typically recorded for 30-40k frames, which corresponds to an acquisition time of 2.5-3.3 minutes. SP-STORM imaging of farred dyes were carried out in a buffer containing 10% (w/v) glucose, 0.5 mg/mL glucose oxidase, 40 µg/mL catalase, and 1% (v/v) beta-mercaptoethanol in 100 mM Tris-HCl buffer (pH 8.0).

### Data analysis

The collected single molecule imaging data were analyzed by either ThunderStorm^22^ ImageJ plugin or Insight3. The single molecules in all three channels were first identified and localized independently. To identify the same single molecule in all channels, a previously developed correlation analysis procedure was adapted.^21^ We first imaged fluorescent beads on coverslip and use their localized positions to build a transformation matrix. The transformation matrix can then be used to project the spatial coordinates of localized single molecule positions from sine- and cosine-modified channel to the refence channel. The projected molecular positions are then compared to those obtained in the refence channel. Molecular positions that are within the same imaging frame and within the localization precision are considered as from the same molecule. Their photon intensities were used for constructing phasor plot (G, S) and calculating the phase angle (ϕ) using the following equations:

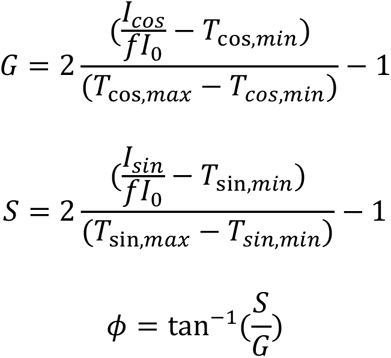

where I_0_, I_sin_, I_cos_ are photon intensity of single molecules in reference, sine, and cosine channels, respectively. f (∼1.17) is the correction factor for the intrinsic difference in transmission efficiency between reference and sine/cosine channels. T_cos,max_ and T_cos,min_ are the maximum and minimum transmission of the cosine filter in the confined spectral window. T_sin,max_ and T_sin,min_ are the maximum and minimum transmission of the sine filter in the confined spectral window. Applying the calibration curve of spectral mean-phase angle relationships, we obtained the spectral mean (λ_mean_). The final localized positions of single molecules were weighted average from those in all channels. Combining the obtained spectral mean and weighted average localized positions of single molecules, hyperspectral dSTORM image was rendered using Insight3 software. For classification of single molecules when the information of all dye molecules in the sample was known, boundary conditions in phasor plot were established from single-color labelled sample and used to separate the different dye molecules with a low crosstalk (Fig 4c, Supplementary Fig.17g). Post-data analysis of localized single molecules was done using MATLAB scripts and Insight3.

### Simulation study

The emission spectra of used dyes in this work are broad. The locations of dyes in the spectral phasor plot depend on both the wavelength and the width of emission spectra, which lays the foundation for spectral demixing in phasor analysis. We use simulation results to demonstrate the working principle of spectral demixing based on phasor analysis and the effects of spectral properties on the performance. We simulated the fluorescent emission spectra at six peak wavelengths (650-725 nm with 15 nm spectral interval) and four peak widths (full width at half maximum, FWHM= ∼2.355σ: 10, 20, 50, and 100 nm) using the normalized Gaussian equation:

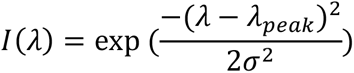

where I(λ), λ_peak_, and σ are intensity at λ, the peak wavelength, and standard deviation of the peak, respectively. The results are shown in Supplementary Fig. 2-3a.

#### Using ideal sine/cosine transmission filters

Spectral phasor analysis using ideal sine/cosine filters were first conducted. The ideal sine/cosine transmission filters are simulated using the following equations:

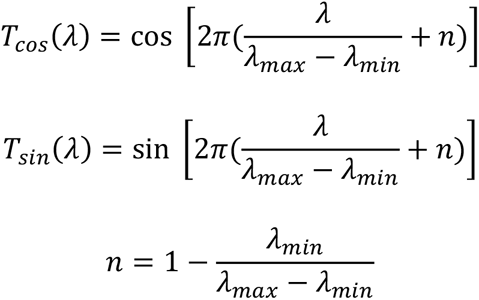

where T_sin_(λ) and T_cos_(λ) are the transmission efficiency for sine and cosine filter at λ; λ_min_ and λ_max_ are the low and high end of detection window of wavelength; n is the correction factor that enables one period of sine/cosine wavefunction within the [λ_min_, λ_max_]. The simulated transmission profiles of ideal sine/cosine filters are shown in Supplementary Fig. 2. Applying the sine/cosine transmission filters on the simulated emission spectra transforms them into two modified emission spectra (Supplementary Fig. 2b-c). The phasor plot (G, S) can then be constructed using the following equation:

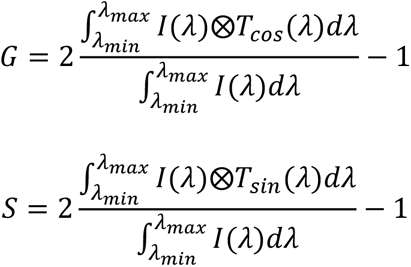

The results in spectra phasor plot for different peak wavelengths and widths are shown in Supplementary Fig. 2d. The results show two obvious trends: larger phase angles for emission at longer wavelengths and smaller phase amplitudes for wider emission spectra. In this work, the wavelength-phase angle relationship is of the most interest. Two types of wavelength-phase angle relationship were plotted using spectral peak and mean respectively (Supplementary Fig. 2e-f). Spectral mean was calculated as the intensity-weighted averaging of wavelength. Significant variation of peak wavelength-phase angle relationships was observed for wide emission spectra (FWHM > 20 nm) due to the confined boundary of spectral detection window. On the contrary, the relationship between spectral mean and phase angle maintains very well (FWHM < 50 nm). In fact, phasor analysis was developed as a non-fitting method for analyzing spectral data and to obtain the spectral mean.

#### Actual sin/cos filters

We further studied the effects of peak wavelength and width in the spectra phasor results when using the actual sine/cosine transmission filters in our experiments. Supplementary Fig. 3 shows the transmission profiles of the sine/cosine transmission filters measured by a UV-Vis spectrometer. Similarly, transformation of emission spectra with actual sine/cosine transmission filters can be done as that used with ideal sine/cosine transmission filters (Supplementary Fig. 3b-c). To construct the phasor plot (G, S), the following modified equations were used to accommodate the min/max transmission of the actual sine/cosine transmission filters:

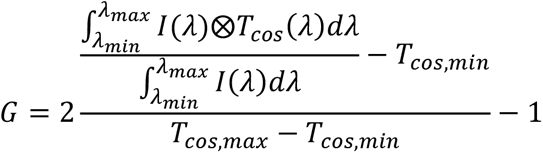

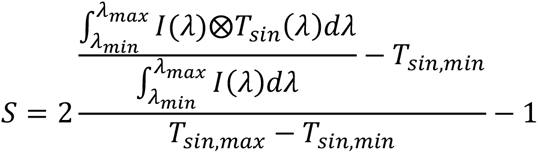

The results are shown in Supplementary Fig. 3d. A nonlinear relationship between wavelength (either spectral peak or spectral mean) and phase angle were obtained (Supplementary Fig. 3e-f), which matches with the calibration results using mechanical slits and fluorescent beads (Fig. 1e). The same results were also observed for the variation of wavelength-phase angle relationships that spectral mean-phase angle relationship maintains the trend for emission spectra with FWHM < 50 nm. In this work, the FWHMs of tested far-red dyes and Nile red dye are less than 50 nm.

## Acknowledgments

This work was supported by the Department of Chemistry at University of Arkansas Fayetteville, and Starter Grant from the Society for Analytical Chemists of Pittsburgh (SACP).

