## Supporting Information for "High Throughput Hyperspectral and Multiplexed Super-Resolution Fluorescence Imaging by SP-STORM"

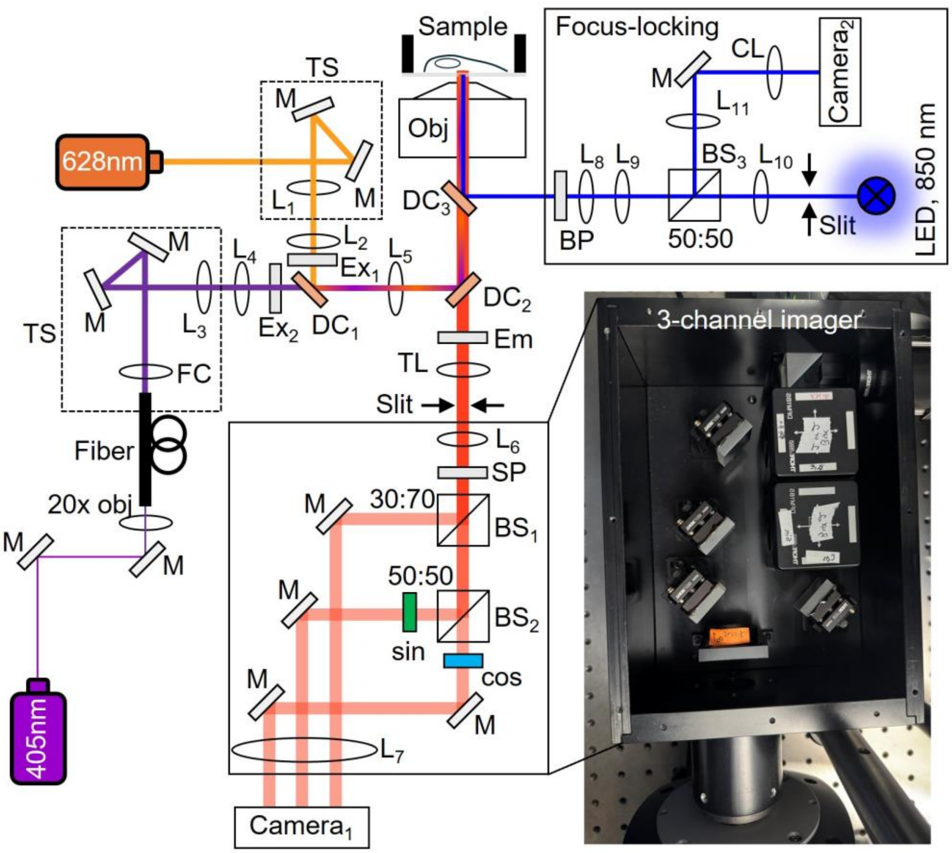


**Supplementary Figure 1. Optical setup of SP-STORM.** The imaging system is built with an Olympus IX81 motorized inverted microscope.

Light sources: Lasers, 100 mW 405 nm (Coherent Obis) and 200 mW 628 nm (MPB communications); LED, 1400 mW 850nm (M850LP1, Thorlabs).

Optics: M, BB1-E02 or BBSQ1-E02 or MRA25-E02 (Thorlabs); Lens1-7, AC254-075-A, AC254-100-A, AC254-100-A, AC254-100-A, AC254-200-A, AC254-125-A, AC254-125-A (Thorlabs); Lens 8-11, LB1945-AB, LB1676-AB, LB1757-AB, LB1901-AB (Thorlabs); CL, LJ1703RM-B (Thorlabs); DC1, 69-204 (Edmund Optics); DC2, RT633rdc (Chroma); DC3, FF750-SDi02 (Semrocks); Ex1, RET633/5x (Chroma); Ex2, ZET405/488/561/640xv2 (Chroma); Em, RET638lp (Chroma); SP, FESH0750 (Thorlabs); BS1, BS019 (Thorlabs); BS2, BS013 (Thorlabs); BS3, BS014 (Thorlabs); BP, FF01-857/30 (Semrock); sine/cosine filters, 700FS80/650FS80 (Andover).

Objectives: 20× air objective (Olympus); 100× Oil TIRF objective (UPLAPO100XOHR, N.A. 1.50, Olympus)

Fiber components: FC, TC18APC-543 (Thorlabs); fiber, P3-S405-FC-1 (Thorlabs).

Mechanic and motorized components: TS, PT1-Z9 and KDC101 (Thorlabs); slit, VA100CP (Thorlabs).

Detectors: camera 1, Andor iXonEM+ Ultra 897; camera 2, DCC1545M (Thorlabs).

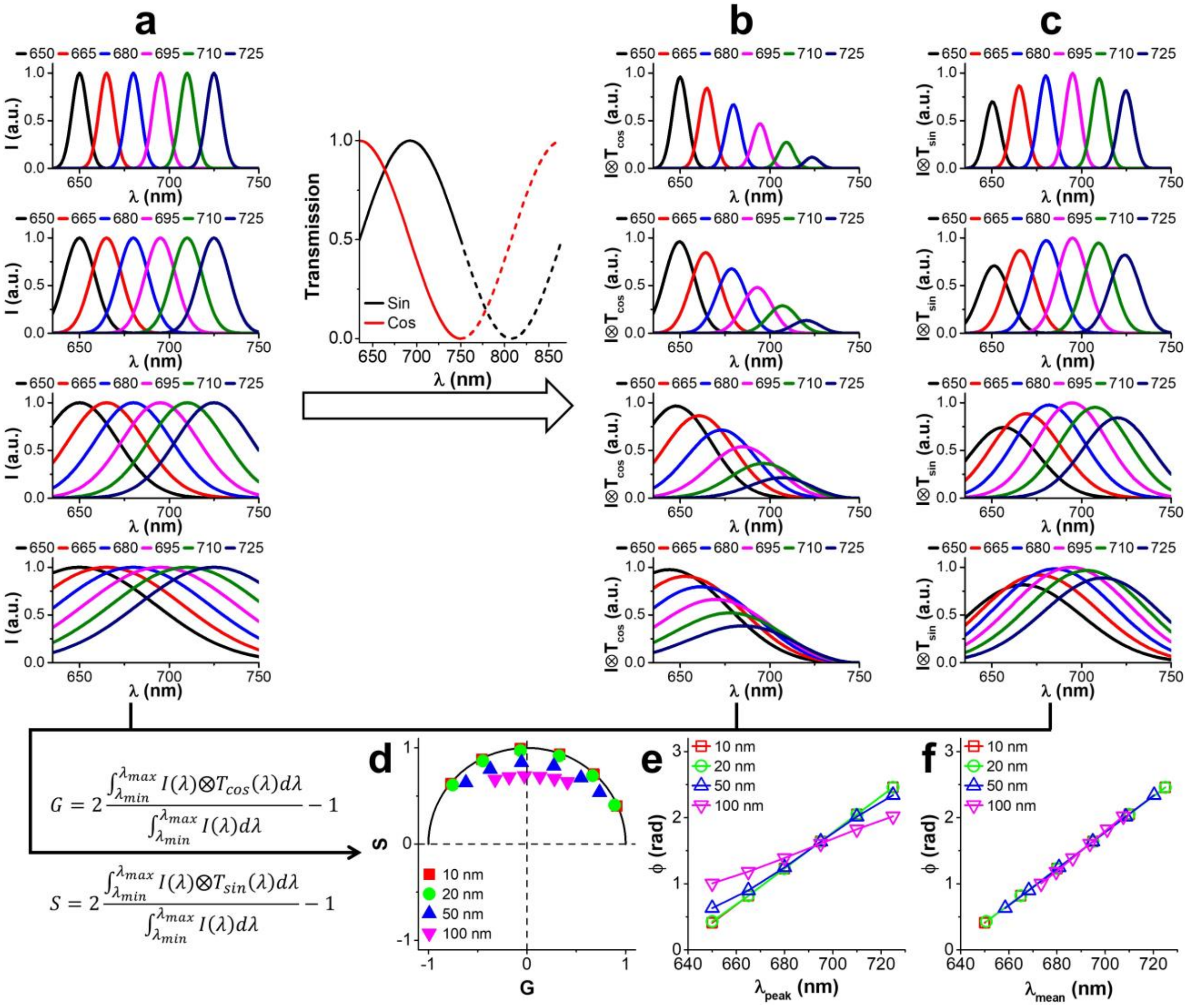


**Supplementary Figure 2. Spectral phasor analysis with ideal sine and cosine transmission filters for far-red dyes.** (a) Simulated emission spectra with Gaussian distribution of different full width of half maximum. (b) Cosine and (c) sine modified emission spectra from (a). (d) Phasor plot. Black solid line denotes results directly calculated from transmission profiles of ideal sine and cosine filters, representing phasor plot at single-wavelength resolution. (e) phase angle-peak wavelength and (f) phase angle-spectral mean relationships, respectively.

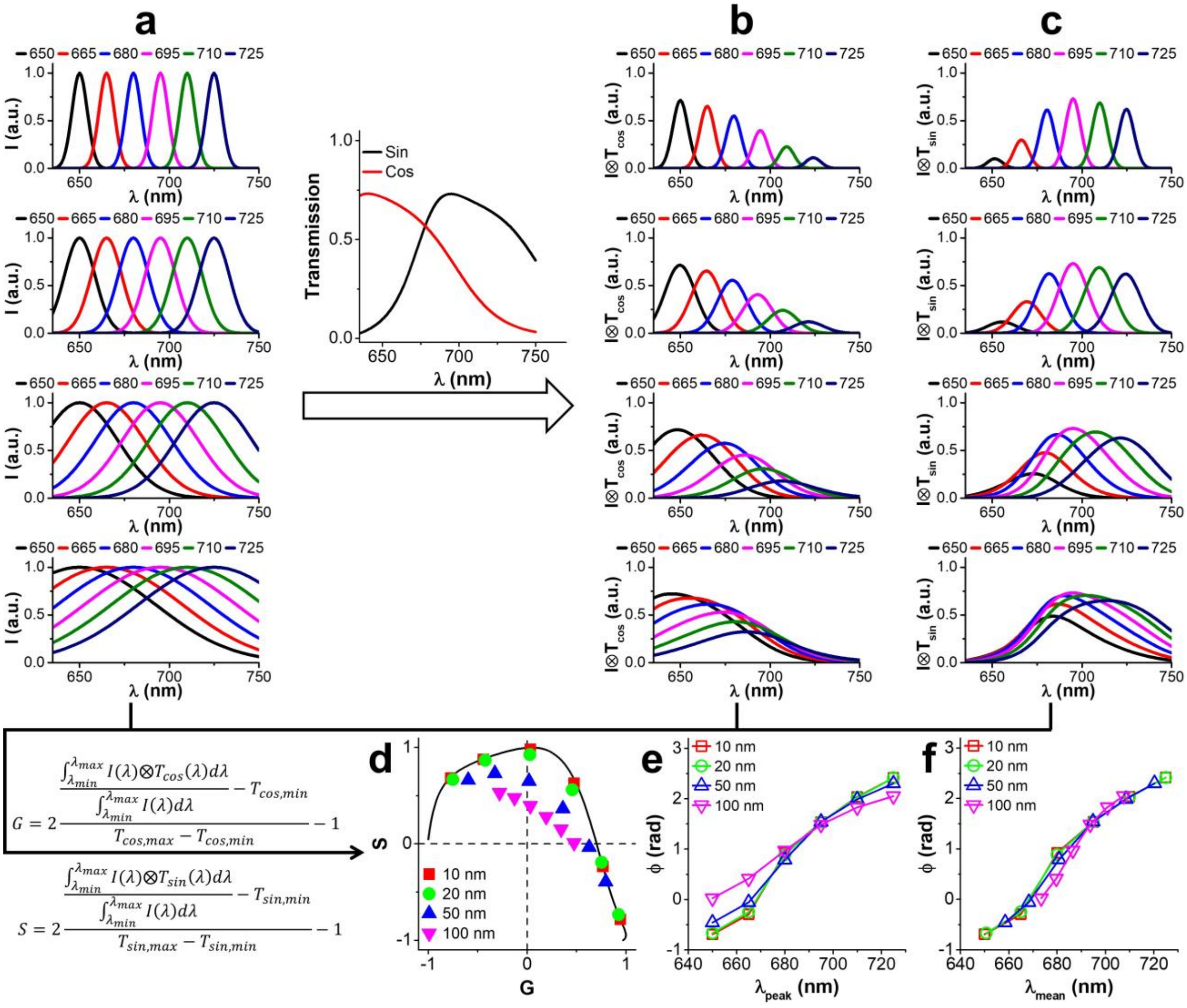


**Supplementary Figure 3. Spectral phasor analysis with actual sine and cosine filters for far-red dyes.** (a) Simulated emission spectra with Gaussian distribution of different full width of half maximum. (b) Cosine and (c) sine modified emission spectra from (a). (d) Phasor plot. Black solid line denotes results directly calculated from transmission profiles of actual sine and cosine filters, representing phasor plot at single-wavelength resolution. (e) phase angle-peak wavelength and (f) phase angle-spectral mean relationships, respectively.

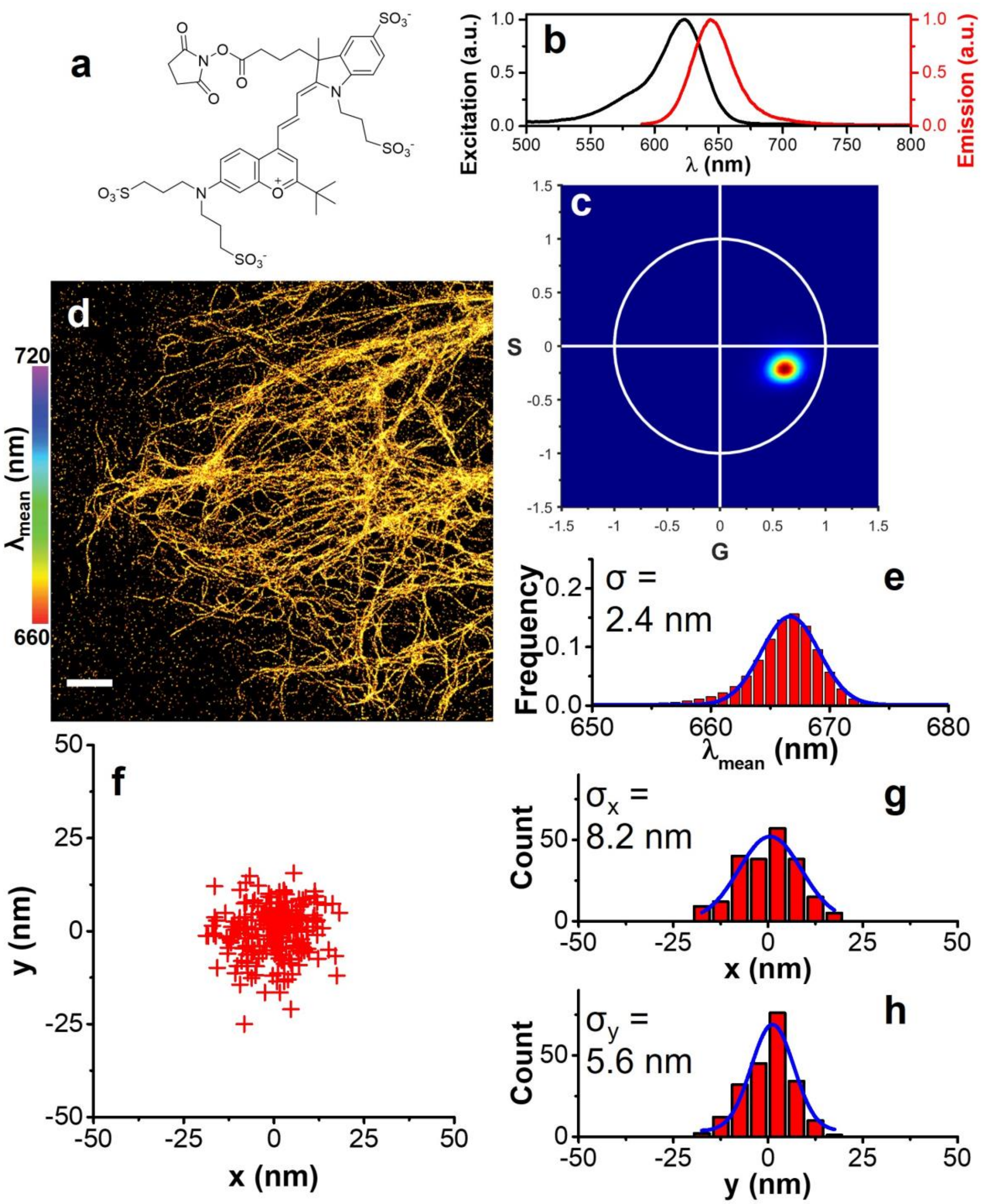


**Supplementary Figure 4. Evaluation of DyLight633 dye for SP-STORM.** (a) Chemical structure. The structure was obtained from the manufacturer. (b) Excitation and emission spectra. (c) Phasor plot of >$10^5$ single DL633 molecules. (d) Hyperspectral dSTORM image of vimentin labelled by DL633 in fixed COS-7 cells. (e) 1D Gaussian fitting the histogram of the spectral mean of single DL633 molecules gives an average of 666.7 ± 2.4 nm (mean ± s.d.). (f) Cluster analysis of locations. (g-h) Fitting histogram distributions in x, y gives standard deviation of $\sigma_x$ = 8.2 and $\sigma_y$ = 5.6 nm respectively. Scale bar: 2 μm.

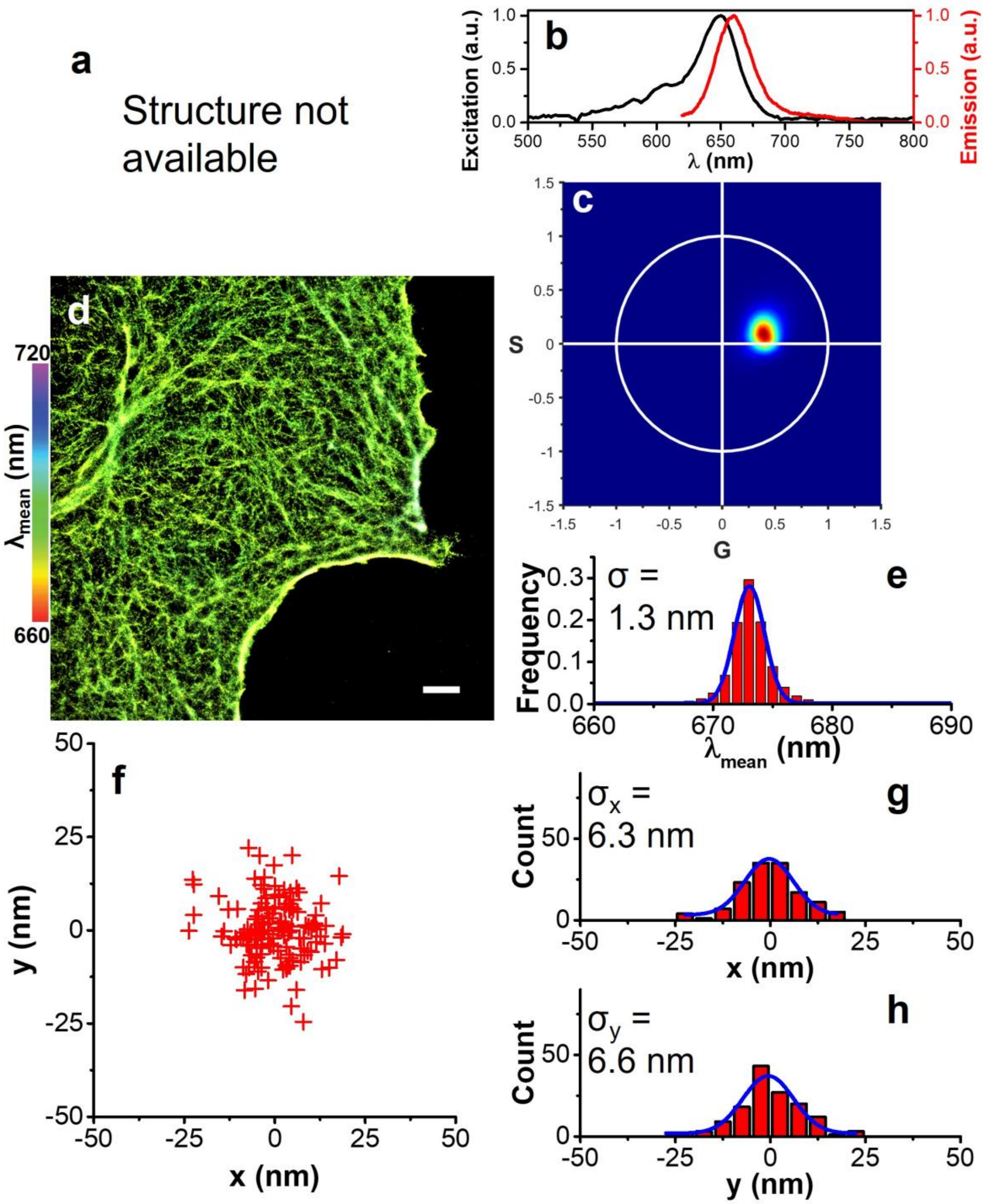


**Supplementary Figure 5. Evaluation of Alexa fluor plus 647 dye for SP-STORM.** (a) Chemical structure is not available. (b) Excitation and emission spectra. (c) Phasor plot of >$10^5$ single AF647$^+$ molecules. (d) Hyperspectral dSTORM image of F-actin labelled by AF647$^+$ in fixed COS-7 cells. (e) 1D Gaussian fitting the histogram of the spectral mean of single AF647$^+$ molecules gives an average of 672.7 ± 1.3 nm (mean ± s.d.). (f) Cluster analysis of locations. (g-h) Fitting histogram distributions in x, y gives standard deviation of $\sigma_x$ = 6.3 and $\sigma_y$ = 6.6 nm respectively. Scale bar: 2 μm.

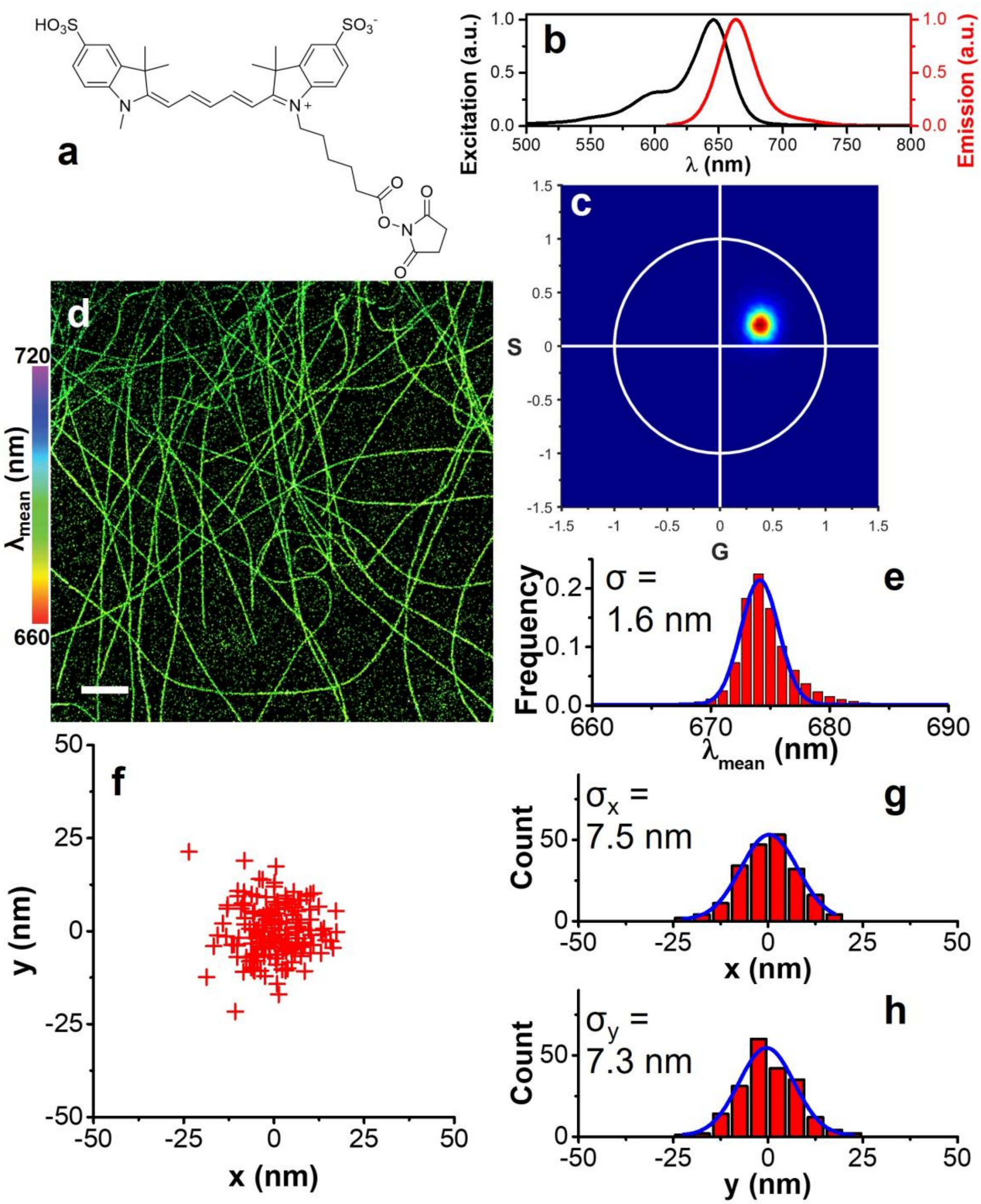


**Supplementary Figure 6. Evaluation of sulfo-cyanine5 dye lumiprobe for SP-STORM.** (a) Chemical structure. The structure was obtained from the manufacturer. (b) Excitation and emission spectra. (c) Phasor plot of >$10^5$ single Cy5 molecules. (d) Hyperspectral dSTORM image of α-tubulin labelled by Cy5 in fixed COS-7 cells. (e) 1D Gaussian fitting the histogram of the spectral mean of single Cy5 molecules gives an average of 674.1 ± 1.6 nm (mean ± s.d.). (f) Cluster analysis of locations. (g-h) Fitting histogram distributions in x, y gives standard deviation of $\sigma_x$ = 7.5 and $\sigma_y$ = 7.3 nm respectively. Scale bar: 2 μm.

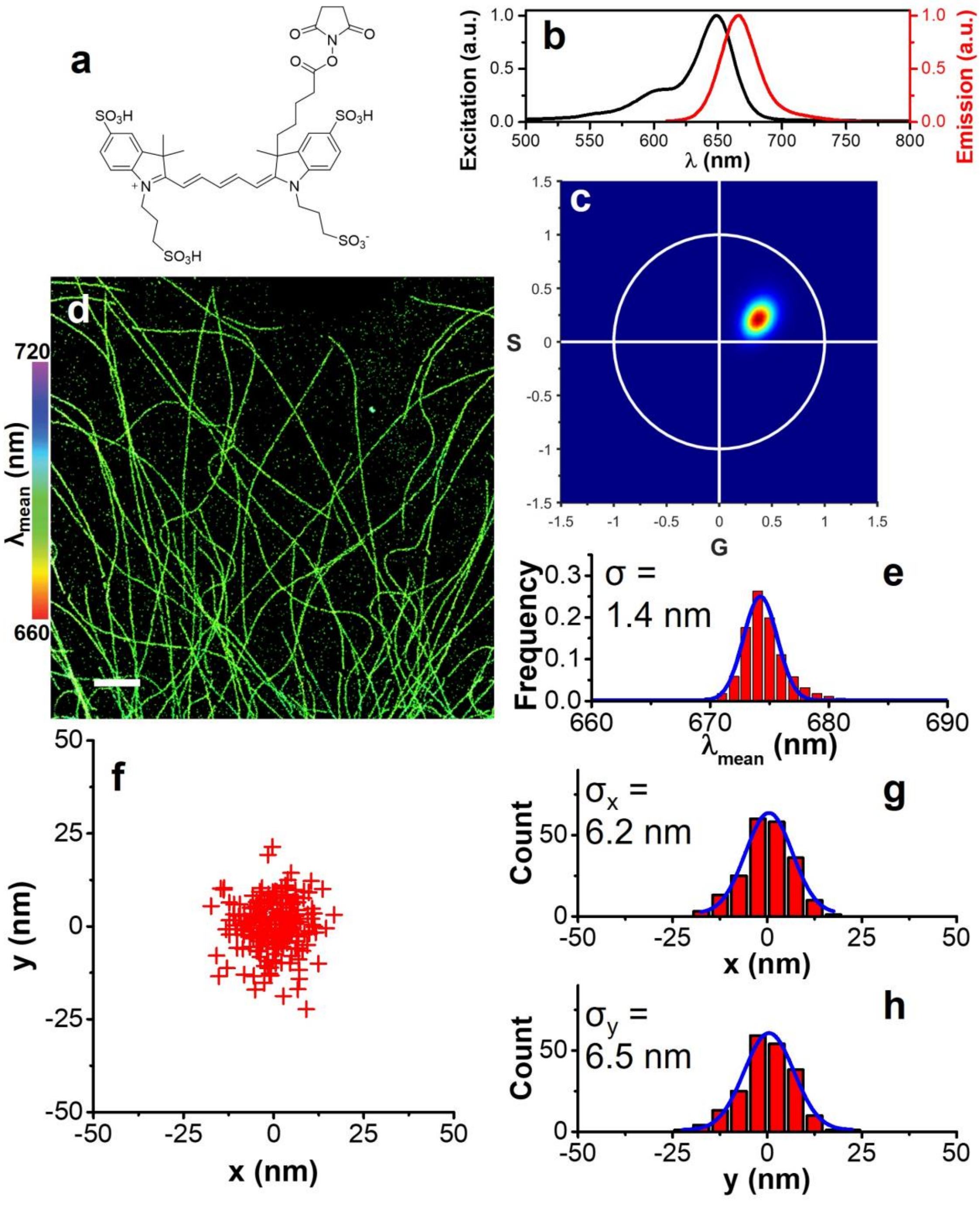


**Supplementary Figure 7. Evaluation of Alexa fluor 647 dye for SP-STORM.** (a) Chemical structure. The structure was obtained from the literature.[1] (b) Excitation and emission spectra. (c) Phasor plot of $>10^5$ single AF647 molecules. (d) Hyperspectral dSTORM image of α-tubulin labelled by AF647 in fixed COS-7 cells. (e) 1D Gaussian fitting the histogram of the spectral mean of single AF647 molecules gives an average of 674.2 ± 1.4 nm (mean ± s.d.). (f) Cluster analysis of locations. (g-h) Fitting histogram distributions in x, y gives standard deviation of $\sigma_x$ = 6.2 and $\sigma_y$ = 6.5 nm respectively. Scale bar: 2 μm.

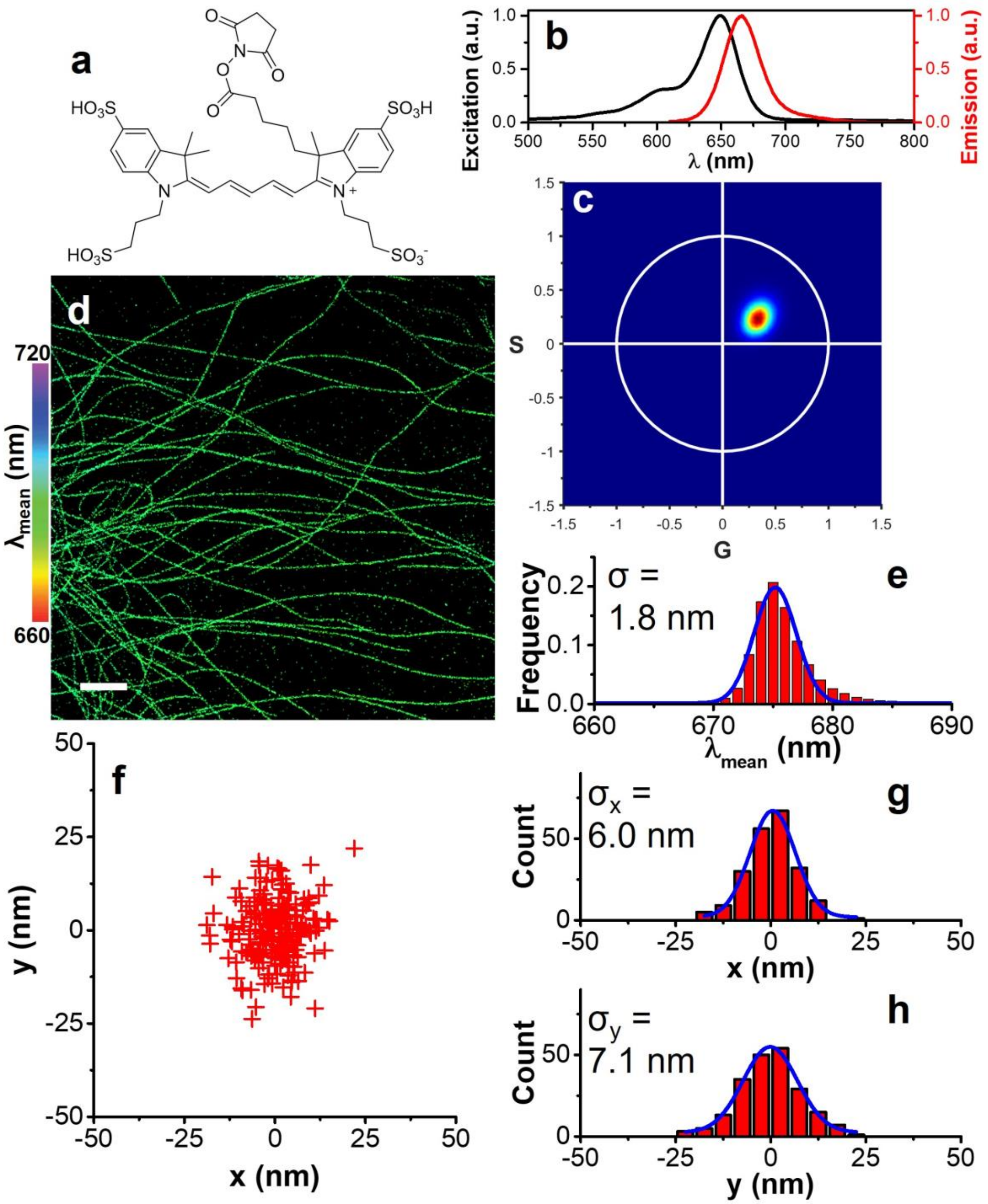


**Supplementary Figure 8. Evaluation of Alexa fluor 647 dye from Lumiprobe for SP-STORM.** (a) Chemical structure. The structure was obtained from the manufacturer. (b) Excitation and emission spectra. (c) Phasor plot of >$10^5$ single $AF647^{lumi}$ molecules. (d) Hyperspectral dSTORM image of α-tubulin labelled by $AF647^{lumi}$ in fixed COS-7 cells. (e) 1D Gaussian fitting the histogram of the spectral mean of single $AF647^{lumi}$ molecules gives an average of 675.2 ± 1.8 nm (mean ± s.d.). (f) Cluster analysis of locations. (g-h) Fitting histogram distributions in x, y gives standard deviation of $\sigma_x$ = 6.0 and $\sigma_y$ = 7.1 nm respectively. Scale bar: 2 μm.

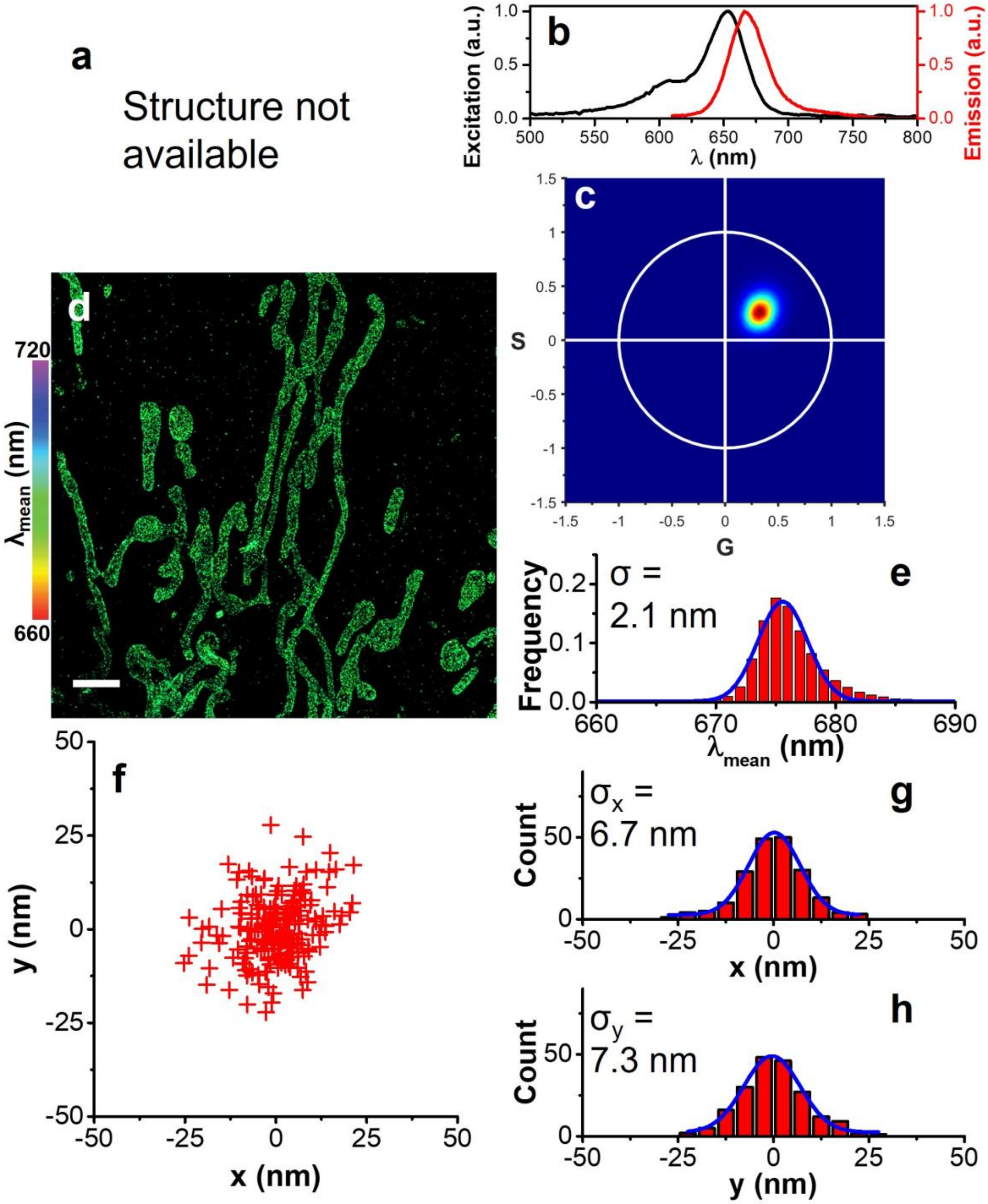


**Supplementary Figure 9. Evaluation of CF647 dye for SP-STORM.** (a) Chemical structure is not available. (b) Excitation and emission spectra. (c) Phasor plot of $>10^5$ single CF647 molecules. (d) Hyperspectral dSTORM image of TOM20 labelled by CF647 in fixed COS-7 cells. (e) 1D Gaussian fitting the histogram of the spectral mean of single CF647 molecules gives an average of 675.6 ± 2.1 nm (mean ± s.d.). (f) Cluster analysis of locations. (g-h) Fitting histogram distributions in x, y gives standard deviation of $\sigma_x$ = 6.7 and $\sigma_y$ = 7.3 nm respectively. Scale bar: 2 μm.

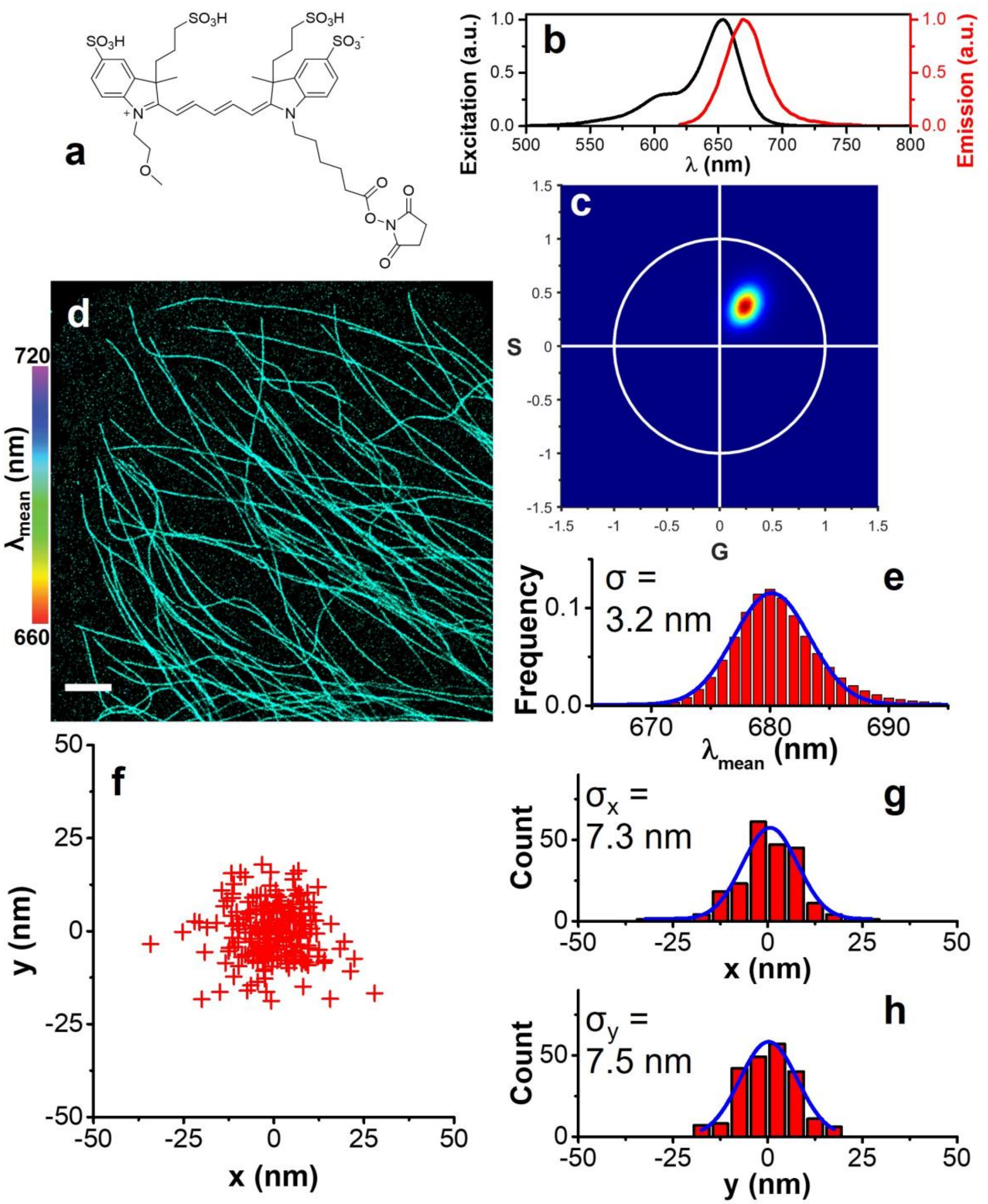


**Supplementary Figure 10. Evaluation of DyLight650 dye for SP-STORM.** (a) Chemical structure. The structure was obtained from the manufacturer. (b) Excitation and emission spectra. (c) Phasor plot of >$10^5$ single DL650 molecules. (d) Hyperspectral dSTORM image of α-tubulin labelled by DL650 in fixed COS-7 cells. (e) 1D Gaussian fitting the histogram of the spectral mean of single DL650 molecules gives an average of 680.1 ± 3.2 nm (mean ± s.d.). (f) Cluster analysis of locations. (g-h) Fitting histogram distributions in x, y gives standard deviation of $\sigma_x$ = 7.3 and $\sigma_y$ = 7.5 nm respectively. Scale bar: 2 μm.

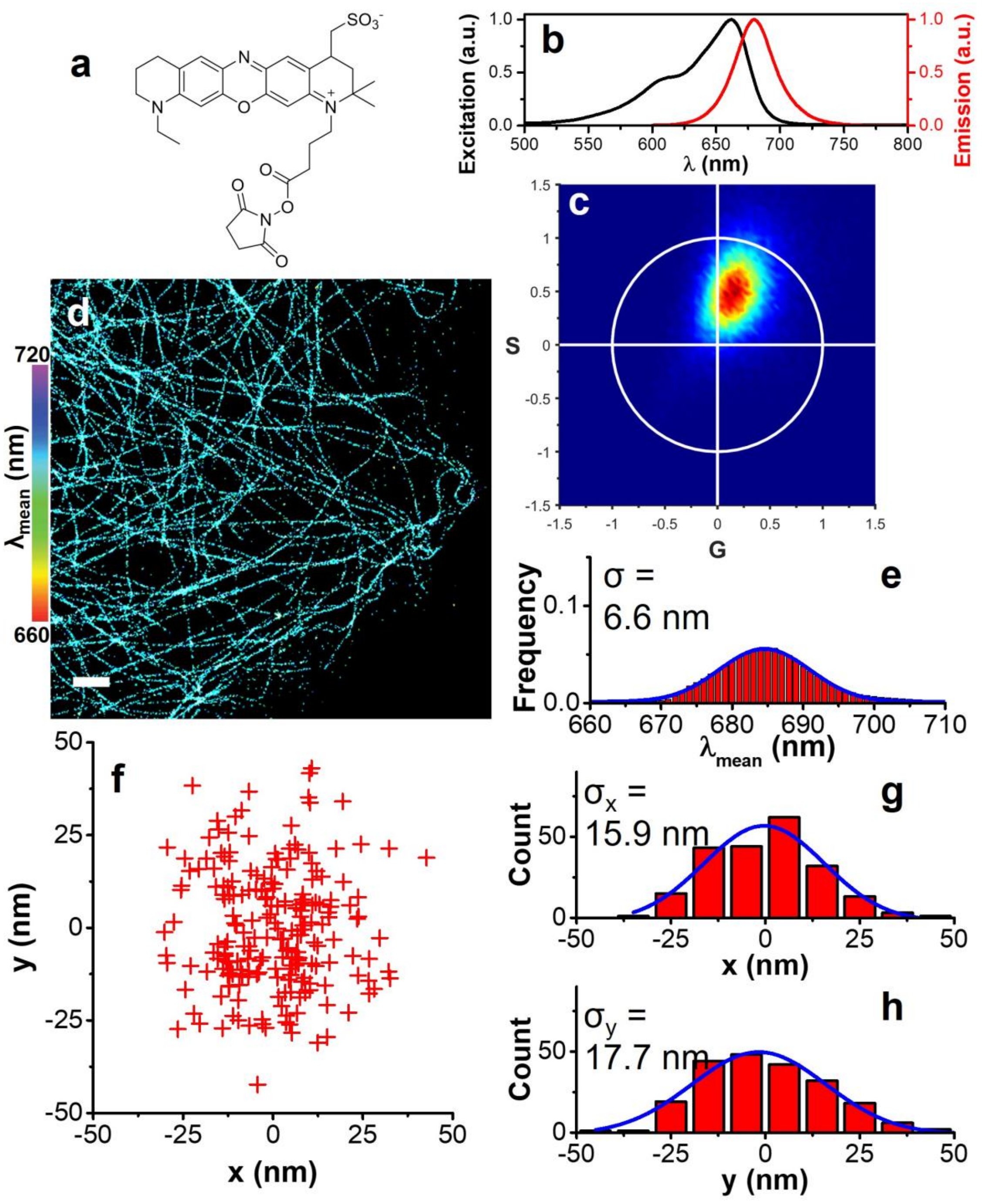


**Supplementary Figure 11. Evaluation of Atto655 dye for SP-STORM.** (a) Chemical structure. The structure was obtained from the literature.[1] (b) Excitation and emission spectra. (c) Phasor plot of >$10^5$ single Atto655 molecules. (d) Hyperspectral dSTORM image of β-tubulin labelled by Atto655 in fixed COS-7 cells. (e) 1D Gaussian fitting the histogram of the spectral mean of single Atto655 molecules gives an average of 684.6 ± 6.6 nm (mean ± s.d.). (f) Cluster analysis of locations. (g-h) Fitting histogram distributions in x, y gives standard deviation of $\sigma_x$ = 15.9 and $\sigma_y$ = 17.7 nm respectively. Scale bar: 2 μm.

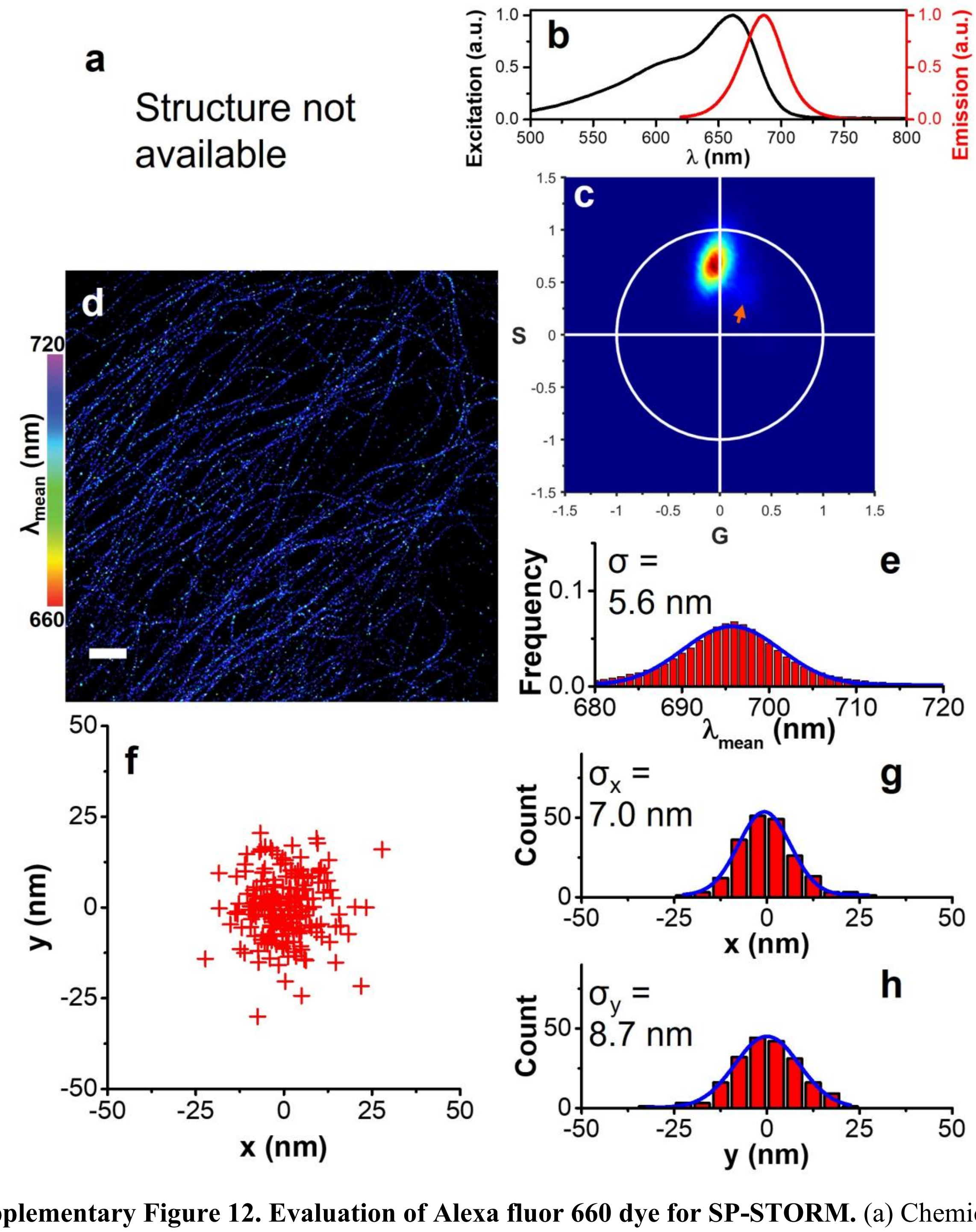


**Supplementary Figure 12. Evaluation of Alexa fluor 660 dye for SP-STORM.** (a) Chemical structure is not available. (b) Excitation and emission spectra. (c) Phasor plot of $>10^5$ single AF660 molecules. A sub-population of smaller phase angles was identified (highlighted by orange arrow). (d) Hyperspectral dSTORM image of β-tubulin labelled by AF660 in fixed COS-7 cells. (e) 1D Gaussian fitting the histogram of the spectral mean of single AF660 molecules gives an average of 695.7 ± 5.6 nm (mean ± s.d.). (f) Cluster analysis of locations. (g-h) Fitting histogram distributions in x, y gives standard deviation of $\sigma_x$ = 7.0 and $\sigma_y$ = 8.7 nm respectively. Scale bar: 2 μm.

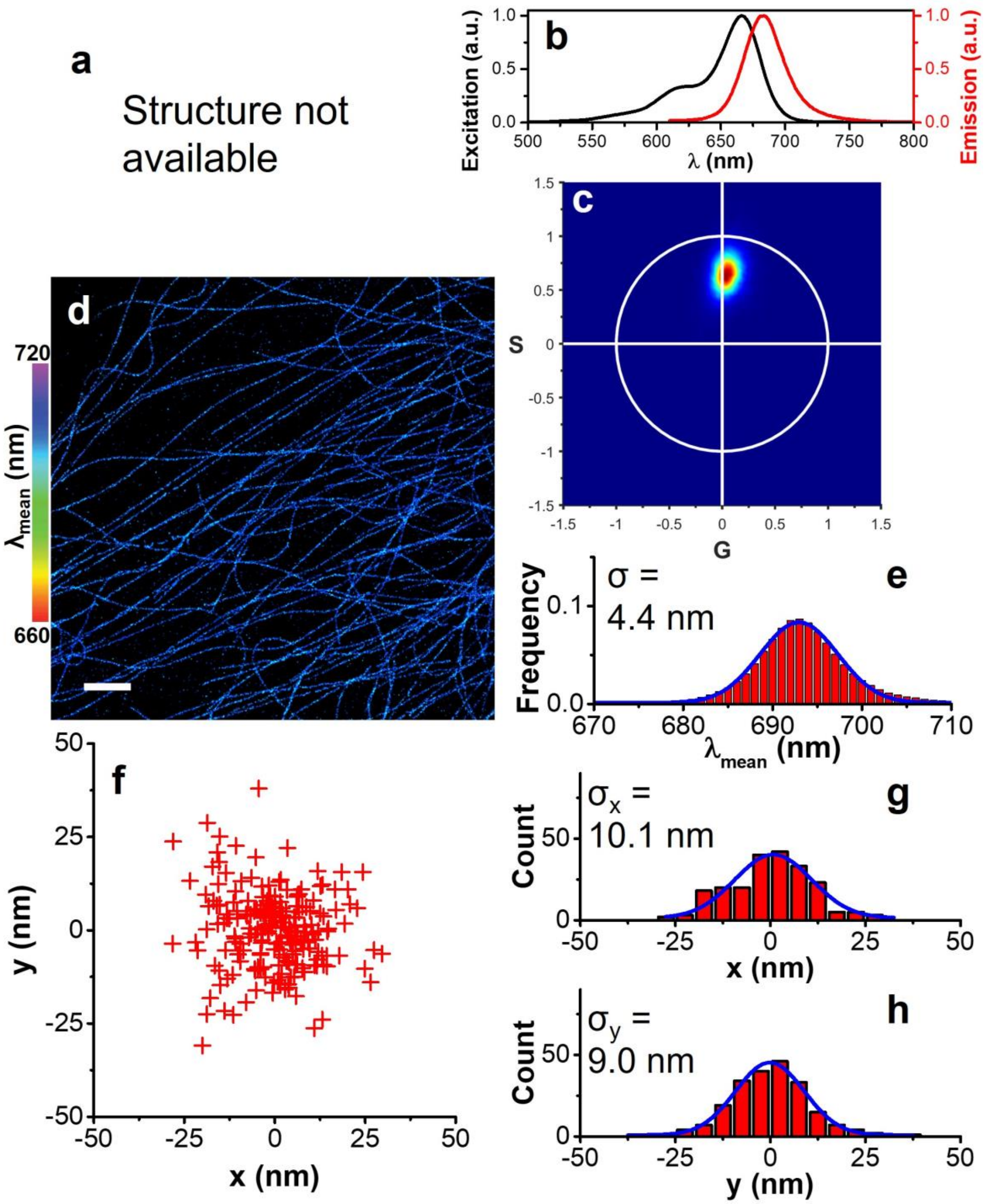


**Supplementary Figure 13. Evaluation of CF660C dye for SP-STORM.** (a) Chemical structure is not available. (b) Excitation and emission spectra. (c) Phasor plot of $>10^5$ single CF660C molecules. (d) Hyperspectral dSTORM image of α-tubulin labelled by CF660C in fixed COS-7 cells. (e) 1D Gaussian fitting the histogram of the spectral mean of single CF660C molecules gives an average of 692.9 ± 4.4 nm (mean ± s.d.). (f) Cluster analysis of locations. (g-h) Fitting histogram distributions in x, y gives standard deviation of $\sigma_x$ = 10.1 and $\sigma_y$ = 9.0 nm respectively. Scale bar: 2 μm.

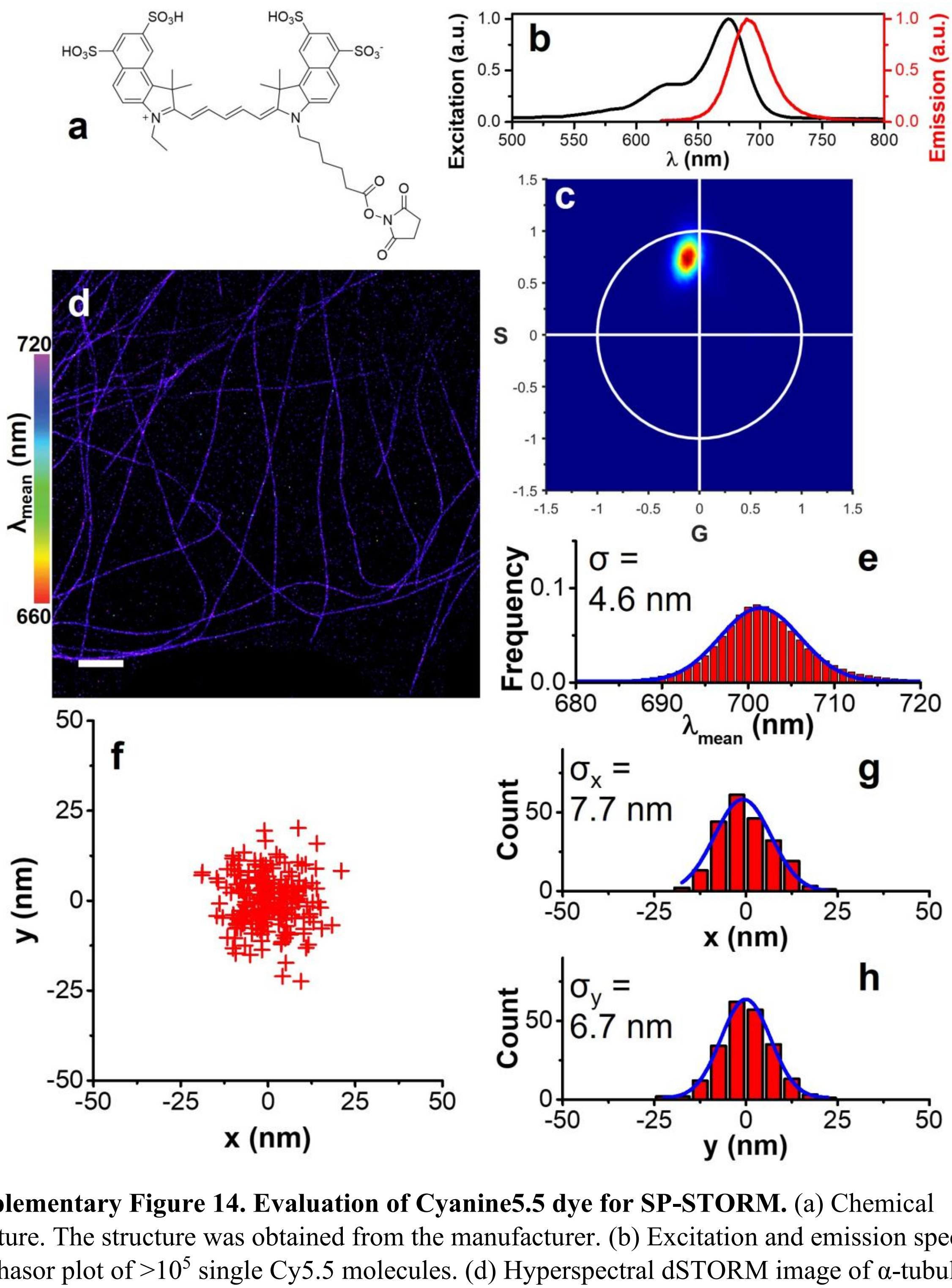


**Supplementary Figure 14. Evaluation of Cyanine5.5 dye for SP-STORM.** (a) Chemical structure. The structure was obtained from the manufacturer. (b) Excitation and emission spectra. (c) Phasor plot of $>10^5$ single Cy5.5 molecules. (d) Hyperspectral dSTORM image of α-tubulin labelled by Cy5.5 in fixed COS-7 cells. (e) 1D Gaussian fitting the histogram of the spectral mean of single Cy5.5 molecules gives an average of 701.4 ± 4.6 nm (mean ± s.d.). (f) Cluster analysis of locations. (g-h) Fitting histogram distributions in x, y gives standard deviation of $\sigma_x$ = 7.7 and $\sigma_y$ = 6.7 nm respectively. Scale bar: 2 μm.

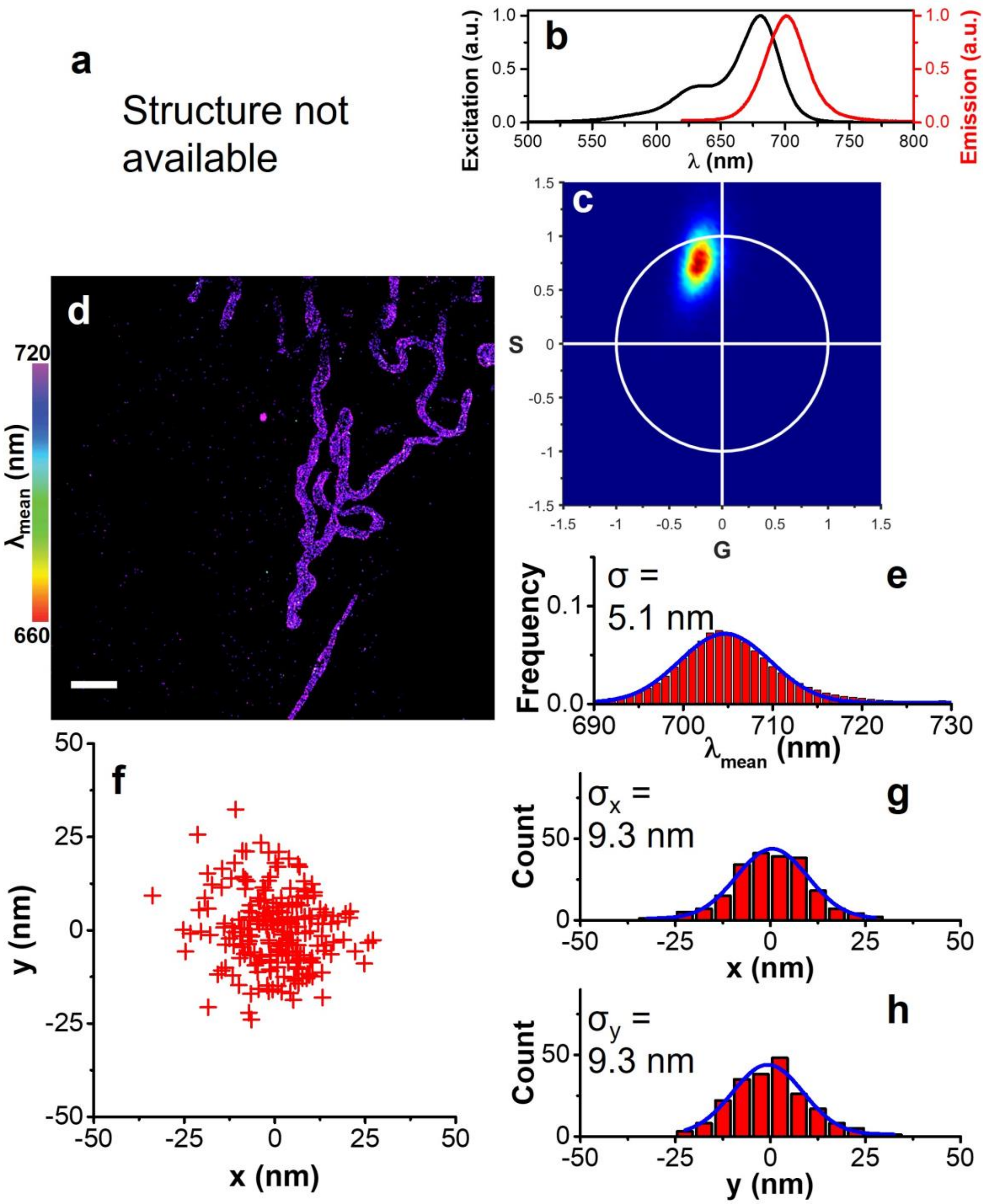


**Supplementary Figure 15. Evaluation of CF680 dye for SP-STORM.** (a) Chemical structure is not available. (b) Excitation and emission spectra. (c) Phasor plot of >$10^5$ single CF680 molecules. (d) Hyperspectral dSTORM image of TOM20 labelled by CF680 in fixed COS-7 cells. (e) 1D Gaussian fitting the histogram of the spectral mean of single CF680 molecules gives an average of 704.6 ± 5.1 nm (mean ± s.d.). (f) Cluster analysis of locations. (g-h) Fitting histogram distributions in x, y gives standard deviation of $\sigma_x$ = 9.3 and $\sigma_y$ = 9.3 nm respectively. Scale bar: 2 μm.

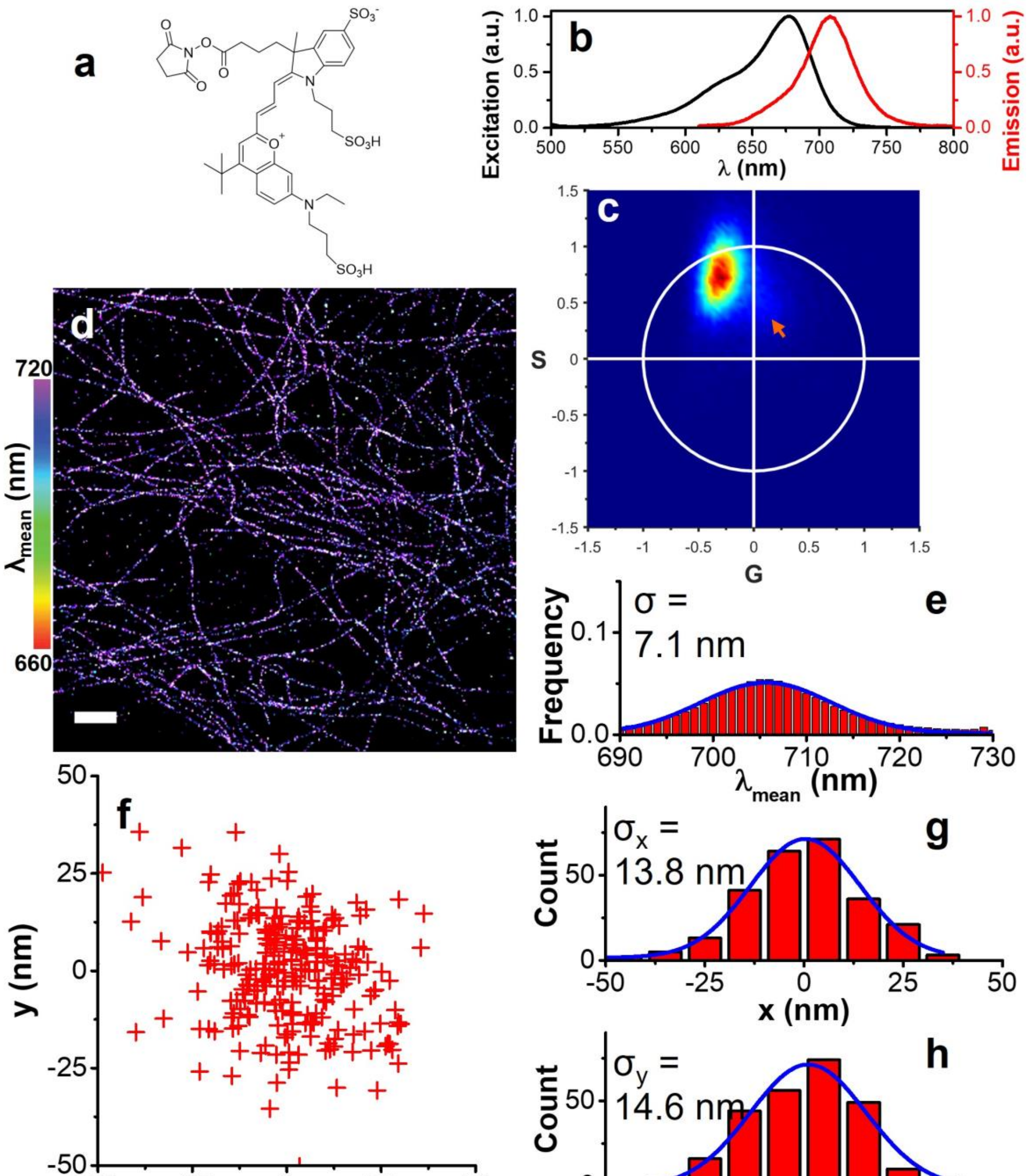


**Supplementary Figure 16. Evaluation of DyLight680 dye for SP-STORM.** (a) Chemical structure. The structure was obtained from the manufacturer. (b) Excitation and emission spectra. (c) Phasor plot of $>10^5$ single DL680 molecules. A sub-population of smaller phase angles was identified (highlighted by orange arrow). (d) Hyperspectral dSTORM image of β-tubulin labelled by DL680 in fixed COS-7 cells. (e) 1D Gaussian fitting the histogram of the spectral mean of single DL680 molecules gives an average of 705.7 ± 7.1 nm (mean ± s.d.). (f) Cluster analysis of locations. (g-h) Fitting histogram distributions in x, y gives standard deviation of $\sigma_x$ = 13.8 and $\sigma_y$ = 14.6 nm respectively. Scale bar: 2 μm.

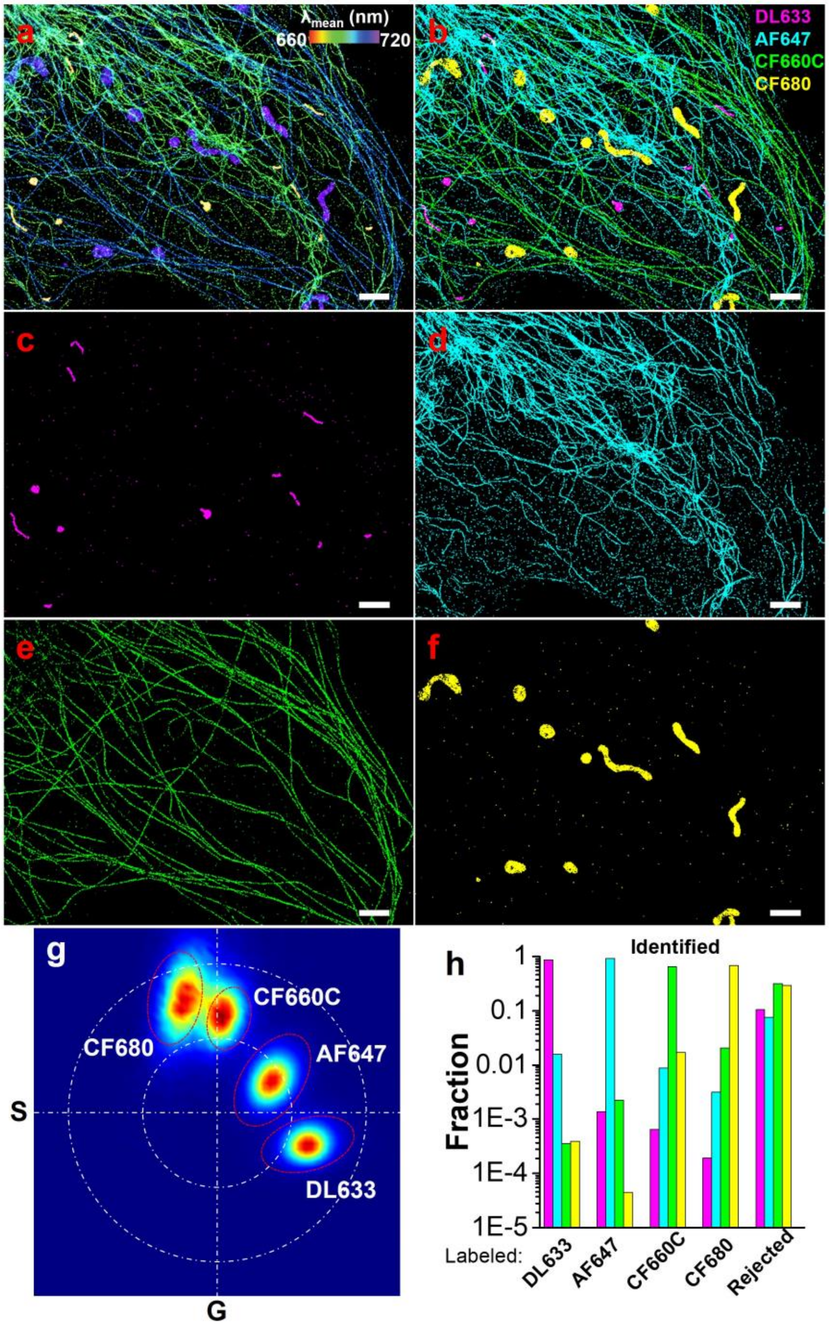


**Supplementary Figure 17. 4-color SP-STORM.** (a) Hyperspectral dSTORM image of four proteins (i.e., PMP70, vimentin, α-tubulin, TOM20) labelled by DL633, AF647, CF660C, and CF680 in fixed COS-7 cells. The color denotes the spectral mean of single molecules. (b-f) 4-color dSTORM images of four proteins after classification of single molecules. (g) Boundary conditions in phasor plot were established from single-color labelled sample and used to separate the different dye molecules. The red dash dot line denotes 1.5-3.0 s.d. of distributions for each dye. The white dash dot line is for illustration location of dyes in the phasor plot with large and small circles as unit and half-unit circle respectively. (h) Color crosstalk between channels. Scale bar: 2 μm.

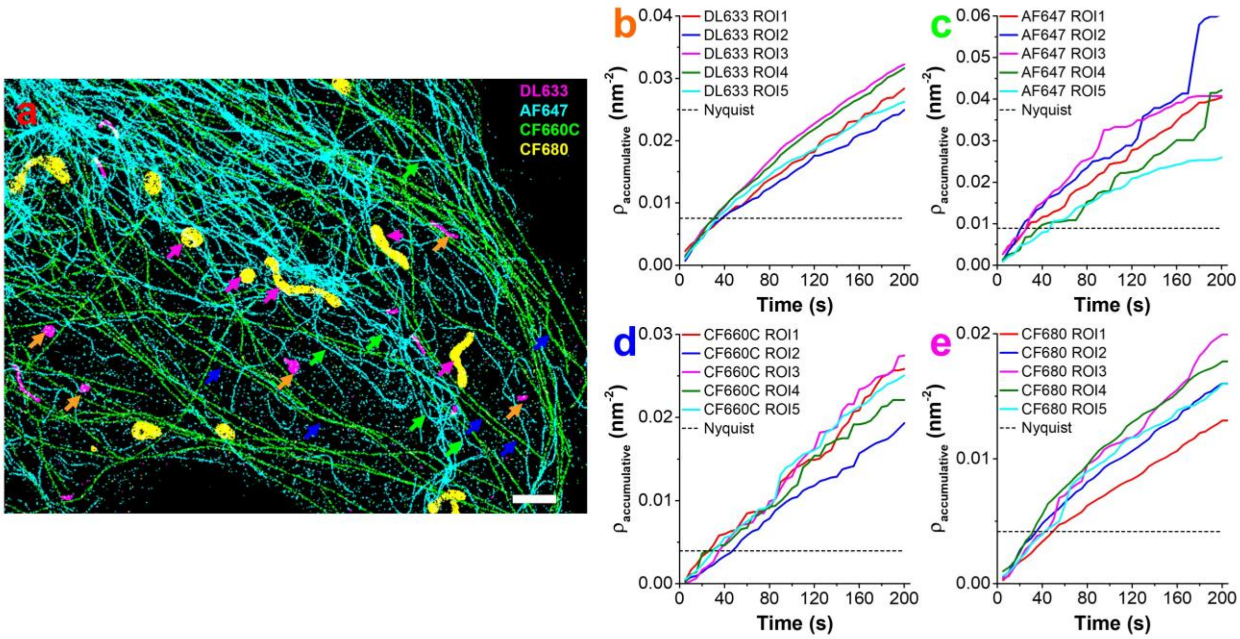


**Supplementary Figure 18. Density of single molecule locations versus imaging acquisition time for 4-color SP-STORM.** (a) dSTORM image of four proteins (i.e., PMP70, vimentin, α-tubulin, TOM20) labelled by DL633, AF647, CF660C, and CF680 in fixed COS-7 cells. (b-e) Density plot of single molecule locations versus time for each dye as highlighted as colored arrows in (a). The short dash line represents minimal Nyquist densities calculated from localization precisions, $\rho = (\frac{2}{2.355\sigma_{xy}})^2$. Scale bar: 2 μm.

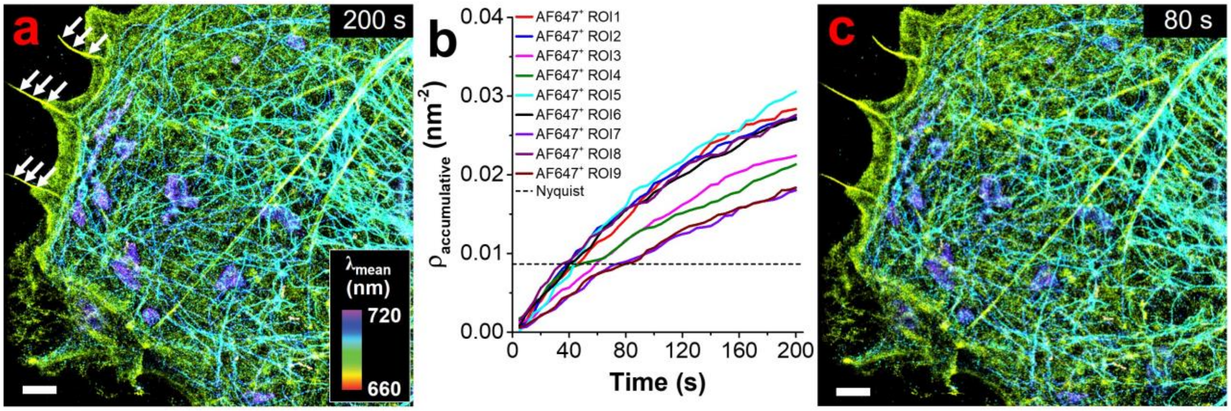


**Supplementary Figure 19. Density of single molecule locations versus imaging acquisition time for 5-color SP-STORM.** (a) Hyperspectral dSTORM image of five proteins (i.e., PMP70, F-actin, vimentin, α-tubulin, TOM20) labelled by DL633, $AF647^+$, DL650, CF660C, and CF680 in fixed COS-7 cells. The color denotes the spectral mean of single molecules. (b) Density plot of single $AF647^+$ molecule locations versus time as highlighted as white arrows in (a). The short dash line represents minimal Nyquist densities calculated from localization precisions, $\rho = (\frac{2}{2.355\sigma_{xy}})^2$. (c) Hyperspectral dSTORM image for a time segment of 80 seconds achieving minimal density based on the Nyquist criterion for $AF647^+$. The subcellular structures of labelled five proteins are already clearly distinguished. Scale bar: 2 μm.
